# Mechanism of CO_2_ and NH_3_ Transport through Human Aquaporin 1: Evidence for Parallel CO_2_ Pathways

**DOI:** 10.1101/2025.02.28.640247

**Authors:** Raif Musa-Aziz, R. Ryan Geyer, Seong-Ki Lee, Fraser J. Moss, Walter F. Boron

## Abstract

The traditional view had been that dissolved gases cross membranes simply by dissolving in and diffusing through membrane lipid. However, some membranes are impermeable to CO_2_ and NH_3_, whereas some aquaporin (AQP) water channels—tetramers with hydrophobic central pores—are permeable to CO_2_, NH_3_, or both. Nevertheless, we understand neither the routes that CO_2_ and NH_3_ take through AQP tetramers, nor the basis of CO_2_/NH_3_ selectivity. Here, we show— for human AQP1 (hAQP1)—that virtually all NH_3_ and H_2_O pass through the hydrophilic, monomeric pores. However CO_2_ passes both through the monomeric pores and another pathway. We expressed hAQP1 in *Xenopus* oocytes and used microelectrodes to monitor the maximal surface-pH transient (ΔpH_S_) caused by CO_2_ or NH_3_ influxes. We found that p-chloromercuribenzene sulfonate (pCMBS)—which reacts with C189 in the monomeric pore—eliminates the entire hAQP1-dependent (*) NH_3_ signal (ΔpH_S_*)_NH3_, but only half of the signals for CO_2_ (ΔpH_S_*)_CO2_ or osmotic water permeability *P*_f_*. 4,4’-diisothiocyanatostilbene-2,2’-disulfonate (DIDS), eliminates the remaining (ΔpH_S_*)_CO2_ but has no effect on (ΔpH_S_*)_NH3_ or *P*_f_*. Together, the two drugs completely eliminate the CO_2_ permeability of hAQP1. When we express hAQP1 in *Pichia pastoris*, treat spheroplasts with DIDS, and examine hAQP1 by SDS-PAGE, reactivity with an anti-DIDS antibody shows that DIDS crosslinks hAQP1 monomers. Our results provide the first evidence that a molecule can move through an AQP via a route other than the monomeric pore, and raise the possibility that selectivity depends on the extent to which CO_2_/NH_3_ move through monomeric pores vs. an alternate pathway (e.g., the central pore).

**Key Points:**

- Some membranes have little or no CO_2_ permeability, absent protein channels like aquaporin-1 (AQP1).
- We confirm that, during CO_2_ influx, heterologous expression of human AQP1 (hAQP1) in *Xenopus* oocytes increases the magnitude of the transient surface-pH increase by an amount (ΔpH_S_*)_CO2_, measured with microelectrodes. During NH_3_ influx, hAQP1 expression increases the magnitude of the transient pH_S_ decrease by (ΔpH_S_*)_NH3_.
- p-chloromercuribenzene sulfonate (pCMBS), which reacts with C189 in the monomeric pore, reduces (ΔpH_S_*)_CO2_ by; (ΔpH_S_*)_NH3_, to zero; and AQP1-dependent osmotic water permeability (*P*_f_*), by half.
- 4,4’-diisothiocyanatostilbene-2,2’-disulfonate (DIDS) reduces (ΔpH_S_*)_CO2_ by half, but has no effect on (ΔpH_S_*)_NH3_ or *P*_f_*. DIDS crosslinks AQP1 monomers expressed in *Pichia pastoris*.
- Together, pCMBS+DIDS reduces (ΔpH_S_*)_CO2_ to zero. The C189S mutation of AQP1 eliminates the effects of pCMBS, but not DIDS. Our results thus show that CO_2_ traverses AQP1 via the monomeric pore plus a novel, DIDS-sensitive route that may be the central pore.

## Introduction

Before the discovery of specific protein-mediated pathways in cell membranes, investigators had believed that many small molecules (e.g., H_2_O, urea, glycerol, lactic acid)—including CO_2_— freely permeate all membranes to the extent that they can dissolve in and then diffusing through the lipid phase. Solubility-diffusion theory (S-DT) embodies this concept (reviewed in Boron, 2010; Michenkova *et al*., 2021). For molecules other than dissolved gases, the generalizability of this notion faded with the discovery of each new membrane channel or transporter, consistent with the idea that cells evolved to achieve tight control over the transmembrane traffic of small molecules. We now believe that the traffic of such molecules depends on some combination of membrane proteins and S-DT (responsible for non-specific leaks).

Regarding CO_2_ traffic across biological membranes, the first inconsistency in the applicability of S-DT was the demonstration that apical membranes (i.e., facing the lumen) of single gastric glands are impermeable to CO_2_ (Waisbren *et al*., 1994). The second was the discovery that the human (h) aquaporin-1 (AQP1) not only conducts H_2_O of course (Preston & Agre, 1991), but also CO_2_ (Nakhoul *et al*., 1998). Soon after Nakhoul’s observation, Cooper and Boron (1998) found that pCMBS inhibits CO_2_ permeability in oocytes expressing wild-type (WT) hAQP1. Prasad *et al*. (1998) then reported that AQP1 purified from human red blood cells (RBCs) and reconstituted into lipids from *E. coli* increases CO_2_ permeability, an effect blocked by HgCl_2_. Moreover, Forster *et al*. (1998) found that 4,4’-diisothiocyanatostilbene-2,2’-disulfonate (DIDS) applied to native human RBCs reduces not only 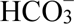 permeability, but also CO_2_ permeability.

Forster and colleagues recognized that an explanation for their data could be that DIDS blocks a protein pathway for CO_2_ diffusion through the RBC membrane. Indeed, Endeward *et al*. (2006) later found that AQP1 is responsible for about half of the CO_2_ traffic across the human RBC membrane, with the rhesus (Rh) complex’s being responsible for most of the rest (Endeward *et al*., 2008), leaving <10% to be accounted for by S-DT. Uehlein *et al*. (2003) reported that NtAQP1 in tobacco plants plays important physiological roles in CO_2_ uptake—driven by a very small air-to-chloroplast gradient—for photosynthesis, stomatal opening, and leaf growth. Wang and colleagues demonstrated that the CO_2_ permeability of tomato aquaporin PIP2;1 is essential for CO_2_-dependent regulation of stomatal aperture (Wang *et al*., 2016).

Besides H_2_O and CO_2_, various AQPs can conduct a variety of substances, including glycerol in the case of members of the aquaglyceroporin subfamily (Gomes *et al*., 2009). Regarding dissolved gases, hAQP1 conducts NH_3_ (Nakhoul *et al*., 2001) and NO (Herrera *et al*., 2006). Moreover, different AQPs can exhibit strong selectivity for CO_2_ over NH_3_ or vice versa (Musa-Aziz *et al*., 2009*a*; Geyer *et al*., 2013). More recent work has suggested that *Nicotiana tabacum* PIP1;3 (Zwiazek *et al*., 2017) as well as hAQP1 (Al-Samir *et al*., 2025) are permeable to O_2_, and that AQP1, the Rh complex, and an unidentified protein are responsibly for nearly all O_2_ permeability in murine RBCs (Moss *et al*., 2025; Occhipinti *et al*., 2025; Zhao *et al*., 2025). These results are consistent with the idea that cells, by regulating channel expression and trafficking to specific membranes, could control the permeability to CO_2_, O_2_, and NH_3_.

Structural studies show that AQP1 is a homotetramer with four independent monomeric pores—lined mainly by hydrophilic residues—that conduct H_2_O (Walz *et al*., 1997; Sui *et al*., 2001). At the core of its square-shaped array of four monomers, AQP1 has a central pore—lined exclusively by hydrophobic residues—with no established function. Each monomer spans the lipid bilayer six times and has cytoplasmic amino and carboxyl termini. As predicted by the “hourglass model” of Preston and Agre (1991), two consensus NPA motifs—on intracellular and extracellular loops that dip in towards the middle of AQP1, near the plane of the membrane—contribute importantly to the monomeric pore. The well-established reduction of osmotic water permeability (*P*_f_) by HgCl_2_ or pCMBS occurs as these agents react with cysteine-189 (C189), two residues upstream from the second NPA (Preston *et al*., 1993). Replacing this cysteine with serine (C189S), which lacks a sulfhydryl group, eliminates this inhibition (Preston *et al*., 1993).

We previously introduced an approach for using blunt, pH-sensitive microelectrodes to monitor transient changes in extracellular-surface pH (pH_S_), caused by the entry of CO_2_ or NH_3_ into *Xenopus* oocytes heterologously expressing membrane proteins (Endeward *et al*., 2006; Musa-Aziz *et al*., 2009*b*, 2009*a*, 2010, 2014*a*). Mathematical modeling has elucidated the interpretation of CO_2_-dependent pH_S_ transients (Somersalo *et al*., 2012; Occhipinti *et al*., 2014; Calvetti *et al*., 2020). Using this pH_S_ approach in a survey of mammalian AQPs, we found that AQPs 1, 6, and 9 are permeable to both CO_2_ (exposure to 5% CO_2_) and NH_3_ (0.5 mM NH_4_Cl); AQP0, the M23 variant of AQP4 (AQP4-M23), and AQP5 are permeable to CO_2_ but not NH_3_; AQPs 3, 7, and 8 are permeable to NH_3_ but not CO_2_; and AQP2 as well as AQP4-M1 appear to be permeable to neither (Musa-Aziz *et al*., 2009*a*; Geyer *et al*., 2013). Using an osmotic-shrinkage assay with an 80-fold higher extracellular [NH_3_] than in our studies, Assentoft et al detected NH_3_ permeability through AQP4-M23 (Assentoft *et al*., 2016). A key unanswered question is how various AQPs exhibit such strikingly different CO_2_/NH_3_ selectivities. The answer presumably lies in the pathways that these solutes take through various AQP tetramers. However, despite insights from molecular-dynamics simulations (Wang *et al*., 2007; Assentoft *et al*., 2016), we still lack physiological evidence that establishes the pathways by which either CO_2_ or NH_3_ moves through any AQP. The purpose of the present study is to explore this problem.

Our approach was to use a combination of cell physiology and biochemistry. Heterologously expressing hAQP1 or its C189S mutant in *Xenopus* oocytes, and using the pH_S_ approach, we assessed the effects of pCMBS and DIDS on CO_2_ and NH_3_ permeation. We also assessed *P*_f_ so that we could normalize CO_2_ and NH_3_ data to H_2_O permeability. We examine pCMBS because mercury—acting on monomeric pores—reduces both the *P*_f_ (Preston *et al*., 1993) and CO_2_ permeability of hAQP1 (Cooper & Boron, 1998), as expressed heterologously in oocytes. We work with DIDS because this drug reduces CO_2_ permeability in RBCs, and in oocytes expressing hAQP1 (Endeward *et al*., 2006). We find that virtually all of the hydrophilic NH_3_ molecules—as well as H_2_O, as expected—pass through the four hAQP1 monomeric pores, which each possess three hydrophilic nodes in their selectivity filters that are located at the center of an otherwise long hydrophobic channel (Sui *et al*., 2001). On the other hand, the more hydrophobic CO_2_ travels both via this pCMBS-sensitive pathway—the only established pathway through any AQP—as well as an independent pathway that we can block with DIDS. Studies on hAQP1 expressed in *Pichia pastoris* indicates that DIDS crosslinks hAQP1 monomers. Our work is the first to demonstrate that a substance can move through an AQP via a pathway—possibly the hydrophobic central pore—that is distinct from the monomeric pore. Considering that mutations in human AQPs are associated with a wide variety of pathological conditions (see (Verkman, 2008; Verkman *et al*., 2008; Wang & Boron, 2025), our work could lead to new insights into disease mechanisms, diagnostic tools, and therapeutic approaches for improving clinical outcomes.

## Methods

For previous summaries of our approach in oocyte experiments, see Musa-Aziz et al. (2010, 2009a, 2009b). The following description is in greater depth.

### Ethical approval and animal procedures

The Institutional Animal Care and Use Committee at Case Western Reserve University approved the protocols for housing and handling of *Xenopus laevis* (approval #2020-0066), used as a source of oocytes, and rabbits (approval # 2010-0146), for generating polyclonal antibodies.

#### Frogs

We purchased adult female *Xenopus laevis* frogs (NASCO Inc., Fort Atkinson, WI, USA) and housed them in a 20-gallon static aquarium, managed by the Animal Resources Center (ARC) of the School of Medicine. For stress mitigation, the tank had six or fewer frogs, and for environmental enrichment, we included a PVC elbow pipe. A charcoal Bio-Bag aquarium power pump (Tetra, Blacksburg, VA) circulated dechlorinated water through the tank. Three times per week, ARC staff fed the frogs with adult *Xenopus* diet (Zeigler Bros. Inc., Gardners, PA), sprinkling the food (10 pellets/frog) into the tank and, after a few hours when the *Xenopus* had fed, removing excess food with a net. As judged necessary, the ARC staff partially changed out water in the tank. Every 90 days, they moved the frogs into a newly cleaned tank that contained, in equal amounts, water from the previous tank and new de-chlorinated water.

We anesthetized frogs by immersion in a solution containing 0.2% tricaine (i.e., MS-222; ethyl 3-aminobenzoate methanesulfonate, catalog # A5040, Sigma-Aldrich, St Louis, MO, USA). When an animal became unresponsive to touch, we removed it from the solution and surgically extracted the ovaries. The animal was euthanized by cardiac excision prior to recovery from anesthesia.

In some experiments, we isolated oocytes from *Xenopus* ovarian lobes shipped overnight from NASCO.

#### Rabbits

We purchased adult female New Zealand white rabbits ^1^ from Charles River Laboratories (Ashland, OH). Following IACUC guidelines, ARC staff obtained a 5-mL pre-immune blood sample from an ear vein, and then inoculated two rabbits—gently restrained in a rabbit-restraint cage, and lightly sedated with intramuscular acepromazine (2 mg/kg)—with fresh DIDS-KLH fusion protein (see below) and Freund’s complete adjuvant via subcutaneous injection on the back of the rabbit. A 10-mL sample of blood (via ear veins, alternating sides) was taken 14– 21 days after the first boost, and the sera were evaluated for antigenicity towards proteins labeled with DIDS. Once a rabbit began producing sufficient titers of polyclonal anti-DIDS antibodies, staff collected 10 mL of blood per kg of body weight (∼30–50 mL) at intervals of 21 to 30 days. After two of such larger blood collections, ARC staff euthanized the animals by cardiac excision and exsanguination, with blood from the terminal collection being transferred to our laboratory.

### Solutions and chemicals

#### OR3 media (for maintaining oocytes)

We follow the protocol described in (Musa-Aziz *et al*., 2010). Briefly, we added to 1.8 L of H_2_O one pack of powdered Leibovitz L-15 media with L-glutamine (catalog # L4386, Sigma-Aldrich), 100 mL of 10,0000 U/mL penicillin/streptomycin solution (cat # 15140122, Thermo Fisher Scientific, Waltham, MA, USA; hereafter abbreviated TFS) and 5 mM HEPES free acid. We then titrated the solution to pH 7.50 with NaOH, periodically measuring osmolality 195 mOsm, and sterile-filtered the solution using a Corning™ Disposable Vacuum Filter/Storage system (catalog # 09-761-107, TFS), and stored it in a cold room for up to 2 weeks before use. Finally, just before use, we sterile-filtered a small volume of solution using a 60-mL syringe (catalog # 14-955-461, TFS) and a sterile in-line Nalgene filter with a 0.22-μm pore size and a 25-mm diameter (catalog # 723-9920, TFS).

#### Solutions for physiology experiments

Table 1 describes the final composition of nine solutions used in the physiology experiments. All were assembled and used at room temperature (∼22°C). Where necessary, we describe any necessary additional steps for correct solution assembly (e.g. the order in which components should be added) below.

**Table 1.**
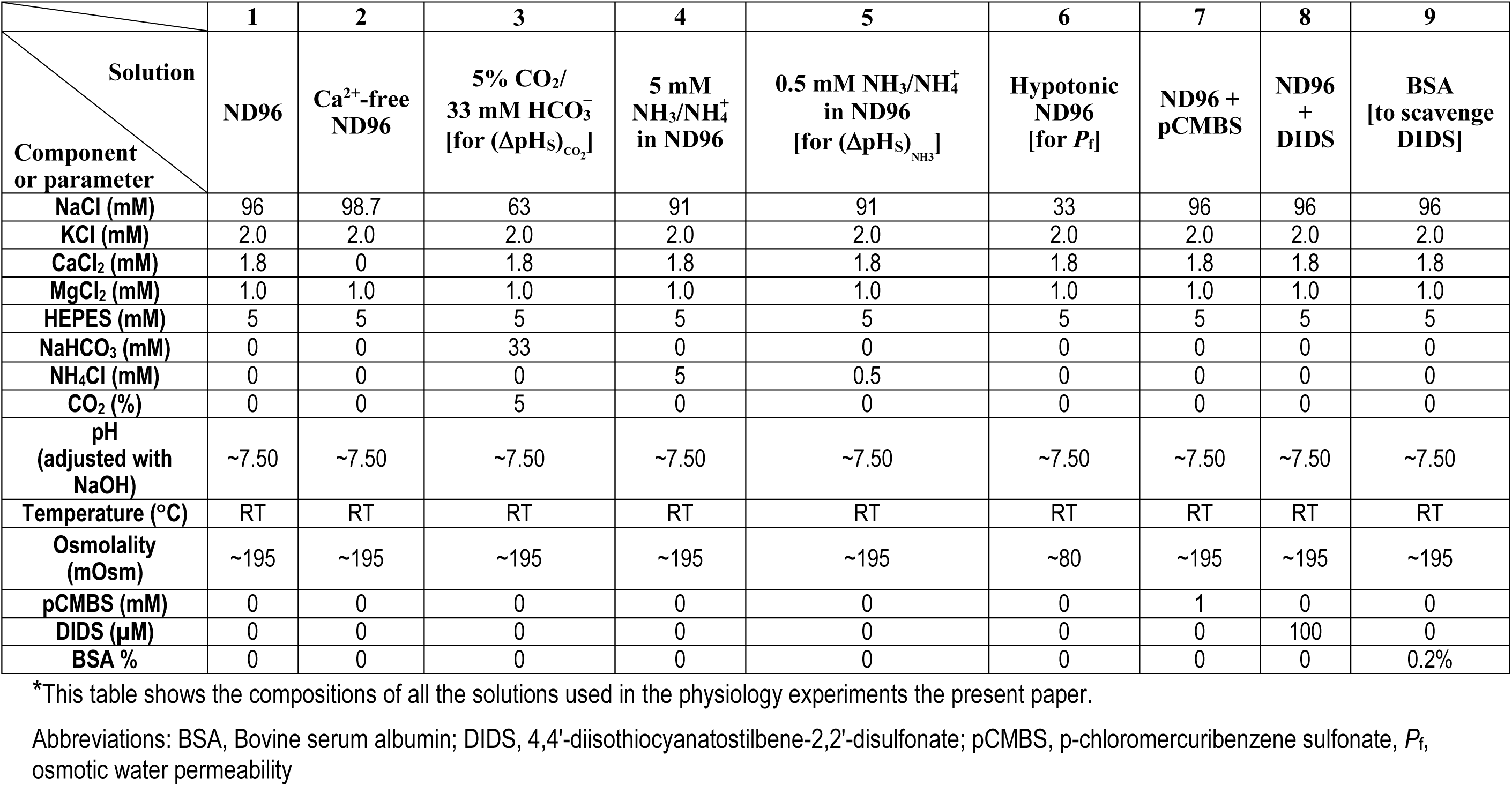
Solutions*.

#### (Solution 3) 5% CO_2_/33 mM 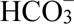 for measuring ΔpH_S_ due CO_2_ influx)

After adding all components except NaHCO_3_, we titrated the solution to pH 7.50, then added the 33 mM NaHCO_3_, and then equilibrated the solution with 5% CO_2_ (balance with air), which brought pH back to 7.50.

#### (Solutions 4 and 5) 0.5 mM NH_3_/NH_4_^+^ in ND96 (for measuring ΔpH_S_ due NH_3_ influx)

We first made the 5 mM NH_3_/NH_4_^+^ in ND96 solution (Solution #4) in which we replaced 5.0 mM NaCl with an equivalent amount of NH_4_Cl, and then diluted this solution 1:10 into ND96.

#### (Solution 6) Hypotonic ND96 (for measuring *P*_f_)

This is a variant of ND96 in which we reduced [NaCl] to lower osmolality to 80 mOsm/kg.

#### (Solution 7) ND96 + pCMBS

In some experiments, we pre-incubated oocytes for 30 min in an ND96 containing 1 mM pCMBS (catalog # C367750, Toronto 196 Research Chemicals, North York, Ontario, Canada), a sulfhydryl reagent, added to solution as dry powder to ND96 (Solution #1).

#### (Solution 8) DIDS

In some experiments, we pre-incubated oocytes for 1 h in an ND96 containing 100 μM DIDS (catalog # D3514, Sigma-Aldrich), an amino-reactive agent added to the solution as a dry powder to ND96 (Solution #1).

#### (Solution 9) BSA (Bovine serum albumin, to scavenge DIDS)

In some experiments, after DIDS pretreatment, we washed off un-reacted DIDS by exposing oocytes to an ND96 to which we added 0.2% BSA (catalog # A9418, Sigma-Aldrich) directly to ND96 (Solution #1).

When assembling all solutions, we measured pH using a Ross electrode (catalog # 927007MD, TFS), a Dual Star pH meter (catalog # 8102BNUWP, TFS), and calibrated with two standards (from TFS), buffer solution pH 6.0 (catalog # SB104-1) and 8.0 (catalog # SB112-20). We measured osmolality with a vapor-pressure osmometer (catalog # 5520 Vapro, Wescor Inc., Logan, UT).

### Oocyte Isolation

We placed ovarian lobes in a sterile 100-mm Petri dish containing ND96, cut them into irregular pieces (<1 cm on an edge) containing ∼10 oocytes each using small iridectomy scissors, poured the mixture into a new Petri dish containing ∼15 mL of Ca^2+^-free ND96, gently agitated the dish on a horizontal shaker^2^ for 10 min, poured off as much liquid as possible, added fresh Ca^2+^-free ND96, and repeated the pour/shake/wash cycle twice more. We then poured off the Ca^2+^-free ND96, before adding ∼15 mL of a freshly-made mixture of Ca^2+^-free ND96 and 2 mg/mL Collagenase Type IA (catalog # C9722, Sigma-Aldrich), and gently agitating the dish on a horizontal shaker for 40 min. We next poured off the collagenase solution, added collagenase-free Ca^2+^-free ND96, gently agitated the dish for 15 min, and repeated the pour/shake/wash cycle twice more. We then poured off the Ca^2+^-free ND96, added ND96 (i.e., containing Ca^2+^), gently agitated the dish on a horizontal shaker for 15 min, and repeated the pour/shake/wash cycle twice more. Finally, we poured off the ND96, added ∼15 mL OR3 media, repeated the pour/wash cycle twice more, poured off most of the liquid, transferred the oocytes and remnant OR3 media to a fresh petri dish containing fresh OR3 media, and incubated the oocytes in an incubator at 18°C. Later the same day, we used a stereomicroscope to sort the oocytes and select individual, defolliculated, mature oocytes (stage V/VI), which we immediately moved to a 6-well plate containing OR3 media (as many as ∼50 oocytes/well), using a fire-sterilized transfer pipette prepared by using a diamond pencil to score the tapered end of a Pasteur pipette at a diameter large enough to accommodate an oocyte, breaking off and discarding the thin end of the pipette, fire-polishing the cut end of the pipette, and attaching the large end of the pipette to a manual, 2-mL Pipette Pump pipetter (catalog # S3-594-3, TFS). We incubated the oocytes overnight at 18°C before injection the following day with cRNA or H_2_O (see <u>below</u>).

### cRNA synthesis

The cDNAs encoding both hAQP1-WT (accession# NM_198098) and AQP1-C189S were gifts of Dr. Peter Agre (Johns Hopkins University). We subcloned the open reading frames of the constructs into the expression plasmid pGH19 (Trudeau *et al*., 1995), a vector containing the 3′ and 5′ untranslated regions of the *Xenopus* laevis β-globin gene. In experiments in which we expressed AQP1-WT or AQP1-C189S in *P. pastoris*—as well as in some control experiments in oocytes—we added an N-terminal (Nt) FLAG-tag (<u>MDYKDDDDKASEFKKKL…</u>; FLAG sequence, underscored; AQP1 sequence, double underscored).

We linearized cDNA constructs using NotI (for untagged AQP1-WT and AQP1-C189S) or XhoI (for the AQP1-FLAG tagged versions). The linearized DNA was then purified using the QIAquick PCR purification kit (catalog # 28104, Qiagen Inc., Valencia, CA). We then synthesized capped RNA (cRNA) using T3 (for untagged constructs; catalog# AM1348, Ambion, Austin, TX, USA) or T7 (for FLAG-tagged constructs; catalog# AM1344, Ambion) mMessage mMachine kits (Ambion, Austin, TX, USA). The cRNA was purified using the RNeasy MinElute RNA Cleanup Kit (catalog # 74204, Qiagen). The cRNA concentration was determined based on ultraviolet absorbance at 260 nm, and quality was assessed according the A260/280 ratio and gel electrophoresis.

### Injection of oocytes with cRNA or water

One day after isolation and defolliculation, we injected stage V–VI oocytes with 25 nL of cRNA (1 ng/nL) encoding AQP1-WT, its C189S mutant, or the FLAG-tagged versions, or ∼25 nL of water as a control. The injection apparatus consisted of a Nanoject II Variable Volume Automatic Injector (Drummond Scientific Company, Broomall, PA) connected to an injection needle that we pulled from 10-μl microdispenser capillary glass tubing (catalog # 3-000-210-G, Drummond), using a Model P-97 Flaming/Brown Micropipette Puller (Sutter Instrument Company, Novato, CA) and then modified by manually breaking the sealed end to yield a tip diameter of ∼50-μm. After injection, we transferred oocytes to a 6-well plate (up to ∼50/well) and incubated in OR3 media and maintained them for 3 days in an incubator at 18 °C before use.

### Electrophysiological recordings

For a schematic representation of the experimental chamber, position of the oocyte, and arrangement of electrodes, see figure 8b in Musa-Aziz *et al*. (2010).

#### Chamber and solution delivery

We placed oocytes into the channel (3 mm wide × 2 mm deep × 30 mm long) of a plastic perfusion chamber. We delivered the physiological solutions—or pH standards for calibrating pH microelectrodes at the outset of an experiment (see next section)— at a constant, total flow of 4 mL/min at room temperature (∼22 °C), using a series of dual-syringe pumps (Model 22, Harvard Apparatus, South Natick, MA), each of which drove two 140-mL plastic syringes (Sherwood Medical, St. Louis, MO) of identical content to deliver two solutions at 2 mL/min each.^3^ The two solutions from each pump flowed through separate lengths of Tygon tubing (5/32-inch (∼4.0 mm) outer diameter, 3/32-inch (∼2.4 mm) inner diameter; Ryan Herco Products Corp., Burbank, CA; Formulation R3603-3; OD 4.8 mm /ID 1.6 mm) to two parallel assemblies of pneumatically operated 5-way valves (Clippard Instrument Laboratory, Cincinnati, OH), daisy-chained in such a way that we could switch among identical pairs of solution lines. The output of the parallel switching assemblies flowed into two lengths of Tygon tubing that carried the solutions to the vicinity of the chamber, where the two lines merged in a “mixing T”, the output of which flowed into one end of the chamber channel. A suction device removed the solution at the opposite end of the channel. See Musa-Aziz *et al*. (2014) for details.

#### Measurement of membrane potential and intracellular pH

As described previously (Musa-Aziz *et al*., 2010, 2014*a*), we impaled the oocyte (with the dark animal pole facing upward) with two microelectrodes, one for measuring membrane potential and the other for measuring intracellular pH. Both microelectrodes, with tip diameters of ∼1 μm, we pulled from thin-walled borosilicate microfilament glass tubing (Catalog # G200TF-4, 2.0 mm OD ×1.56 mm ID, Warner Instruments Corporation, Hamden, CT), using the aforementioned P-97 microelectrode puller.

The microelectrodes for measuring membrane potential (*V*_m_) we filled with 3 M KCl. We inserted an electrode into a half-cell/microelectrode holder (model # ESW-F20N, Warner Instruments Corp.), attached the holder to the *V*_m_ probe of an OC-725 two-electrode Oocyte Clamp (Warner Instruments Corp.), and mounted the probe on a model MM-33R micromanipulator (Warner Instruments Corp.). The *V*_m_ microelectrodes had resistances of ∼0.3 – ∼0.6 MΩ.

The microelectrodes for measuring intracellular pH (pH_i_) were identical to the *V*_m_ electrodes except for the following fabrication process. In order to remove moisture, we placed the pulled micropipettes, with tips upwards, into the cylindrical holes of a reusable aluminum block (machined to create “legs” and thereby allow gases to circulate from beneath, and with smaller concentric cylindrical holes extending the entire height of the block to allow access of gases from below, to the interior of the glass capillary), placed the block into the bottom of a 100-mm Pyrex petri dish, covered the electrodes and block with an inverted 250-mL beaker, placed the petri dish on an open metal rack of an oven at 200 °C, and then baked overnight. With the microelectrodes still in the 200 °C oven, we then silanized them by tilting the beaker slightly and depositing 80 μL of *bis*-di-(methylamino)-dimethylsilane (Sigma-Aldrich, catalog #14755) between the legs of the aluminum block, releasing the breaker, and allowing the silane fumes to interact with the microelectrodes for 40 min before removing beaker. The silanized electrodes cured in the oven at 200 °C until ready for use. We used a hand-drawn, soft-glass pipette with a long tip to backfill each pH_i_ electrode with a liquid pH-sensitive sensor (catalog #95293, Hydrogen Ionophore I, mixture B, Fluka Chemical Corp., Ronkonkoma, NY), as described by (Ammann *et al*., 1981), creating a column that extended ∼1 mm from the microelectrode tip. We then backfilled an microelectrode with a buffer solution (containing, in mM, 40 KH_2_PO_4_, 23 NaOH, 15 NaCl, adjusted to pH 7.0), inserted the electrode into a half-cell/microelectrode holder (model # ESW-F20N, Warner Instruments Corporation), attached the holder to one probe of a model FD223 dual high-impedance electrometer, World Precision Instruments [WPI], Inc., Sarasota, FL), and mounted the probe on a model MM-33R micromanipulator (Warner Instruments Corp.). The pH_i_ microelectrodes had resistances of ∼0.3 – 0.6 MΩ.

After placing the tips of the *V*_m_ and pH_i_ microelectrodes in ND96 solution in the chamber channel (which we now refer to as the “bath”), we obtained *V*_m_ by an analog electronic subtraction of the system ground of the OC-725 amplifier from the signal of the *V*_m_ electrode. We similarly obtained the voltage due to pH_i_ by electronically subtracting the signal of the *V*_m_ electrode from the signal of the pH_i_ electrode. For a schematic representation of the configuration of the electronics for obtaining *V*_m_ (this section) and the voltages due to pH_i_ (this section) and pH_S_ (next section), see figure 7c in Musa-Aziz *et al*. (2010). The device that performed the subtractions (Yale University Subtraction Amplifier, v3.1) also appropriately scaled the voltages for the inputs of an analog-to-digital converter, installed in a Windows-based computer. With the tips of the *V*_m_ and pH_i_ electrodes (and also the pH_S_ electrode, simultaneously; see next section) in the bath, we obtained the slope of the pH_i_ electrodes as we used the aforementioned solution-delivery system (see previous section) to flow a pH standard at pH 6.0 (catalog # SB104-1) continuously through the chamber channel, recorded the voltage due to that pH, used the valve system to switch the flowing solution to a second, pH-8.0 standard (catalog # SB112-20), and then recorded the corresponding voltage. We typically switched between standards in a pH-6→8→6→8 sequence, accepting pH_i_ microelectrodes that completed their electrical response—which also included the time for the solution in the chamber channel to fully change composition—within ∼5 s (∼2 s for pH_S_ electrodes) and had slopes in the range of 53 to 60 mV/pH unit. After the recordings in the final pH-calibration solution, we re-introduced the ND96 solution into the bath, and then added the oocyte to the chamber channel. After impaling the oocyte with the *V*_m_ electrode, and with the tip of the pH_i_ electrode near the oocyte, we obtained a single-point pH calibration of the pH_i_ microelectrode in the ND96 solution (defined as having a pH 7.50). We then impaled the oocyte with the pH_i_ microelectrode. After stabilization, oocytes had spontaneous *V*_m_ values at least as negative as –40 mV. We continuously recorded pH_i_ to judge the integrity of the oocyte and compare the pH_i_ time course with that of surface pH. In the present paper, we are not presenting the pH_i_ data, which nevertheless are similar to those of our previous studies (Musa-Aziz *et al*., 2009*a*, 2010, 2014*a*, 2014*b*).

#### Measurement of surface pH

We measured surface pH with liquid-sensor pH microelectrodes that we fabricated and employed in the same way as we did for pH_i_ microelectrodes (see above), but with four differences. First, we pulled the pH_S_ microelectrodes from standard-walled (rather than thin-walled) borosilicate microfilament glass tubing (Part No. G200F-4, 2.0 mm OD ×1.16 mm ID, Warner Instruments). Second, we used a microforge to break off and fire polish the tips (inner diameter ≅ 20 μm at tip) as one would for a giant-patch pipette (Musa-Aziz *et al*., 2010). It was to facilitate the fire polishing that we used the standard-walled glass tubing. Third, we used a vented microelectrode holder (model # ESW-F20N, Warner Instruments Corp.) mounted on a model MM-33L micromanipulator. The vent prevented pressure buildup that would otherwise have pushed the liquid pH sensor out of the wide electrode tip as we pushed the pH_S_ electrode into the holder. Note that we attached the microelectrode holder of the pH_S_ electrode to the second of two inputs of the FD223 electrometer. And fourth, after obtaining the pH_S_-electrode slope (which we did at the same time as obtaining the pH_i_ electrode slope, as described above) and single-point calibration of the pH_S_ electrode in ND96, we used an ultra-fine computer-controlled micromanipulator (model MPC-200 system, Sutter Instrument Company) to positioned the blunt tip of the pH_S_ electrode near the oocyte’s equator (i.e., between the upward-facing animal pole and downward-facing vegetal pole), and ∼5 degrees behind the meridian (i.e., barely in the “shadow” of the flowing extracellular solution) until the pH_S_ electrode tip just touched the surface of the oocyte (Musa-Aziz *et al*., 2010, 2014*a*). We then further advanced the tip by ∼40 μm to create a slight dimple in the membrane. Periodically during the experiment (indicated by vertical gray bands in each panel of Figure 2*b*, Figure 3, and Figure 6), we withdrew the electrode ∼300 μm from the oocyte for recalibration in the bulk extracellular fluid (bECF) of the bath (i.e., pH 7.50).

The external reference electrode for the pH_S_ measurement was a calomel half-cell, bridged to the chamber channel via a long glass micropipette filled with 3M KCl, connected to a model 750 electrometer (WPI), and positioned so that its tip was just downstream of the oocyte. We obtained the pH_S_ signal by an analog electronic subtraction of the calomel signal from the signal of the pH_S_ electrode.

We established a virtual ground with an Ag/AgCl half cell (connected to the *I*_Sense_ input of the voltage-clamp amplifier) bridged to the chamber by a second glass microelectrode filled with 3M KCl, and positioned the tip of this second bridging pipette close to that of the first, that is, just downstream of the oocyte.

#### Measurement of osmotic water permeability

We measured *P*_f_ as described by (Zhang *et al*., 1990; Preston *et al*., 1992, 1993). Using an approach similar to those in previous studies by our group (Virkki *et al*., 2001; Musa-Aziz *et al*., 2009*a*), we dropped an oocyte into a Petri dish containing a hypotonic solution (80 mOsm; Solution #6, Table 1) to induce cell swelling and, while illuminating from beneath the dish, used a video camera to obtain images of the oocyte (1/s × 60 s). Using a small metal sphere next to the oocyte as a size reference, we determined the projection areas of the oocyte, and—assuming the oocyte to be a perfect sphere—computed idealized oocyte volume (V_Oocyte_) and idealized oocyte surface area as a function of time. We estimated the actual surface area (*S*, in cm^2^) by multiplying the idealized surface area by a factor of 9 (Chandy *et al*., 1997). We computed *P*_f_ from the following equation:

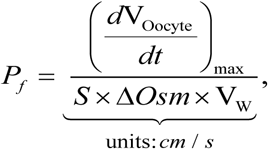

the form of which differs slightly from that of Zhang *et al*., (1990).^4^ Here, (*d*V_Oocyte_/*dt*)_max_ is the maximum time rate of change of cell volume (i.e., the maximal H_2_O influx, or *J*_V,max_, in cm^3^ s^−1^), Δ*Osm* is the initial osmotic gradient across the oocyte membrane (i.e., 195 – 80 = 115 mOsm, expressed as mol cm^−3^), and V_w_ is the molar volume of water (18.1 cm³ mol^−1^). As for the electrophysiological studies, we performed all *P*_f_ assays at room temperature (∼22 °C).

### Protocols for physiological experiments

Figure 1*A–J* illustrates the sequence of events in our experimental protocols.

**Figure 1:**
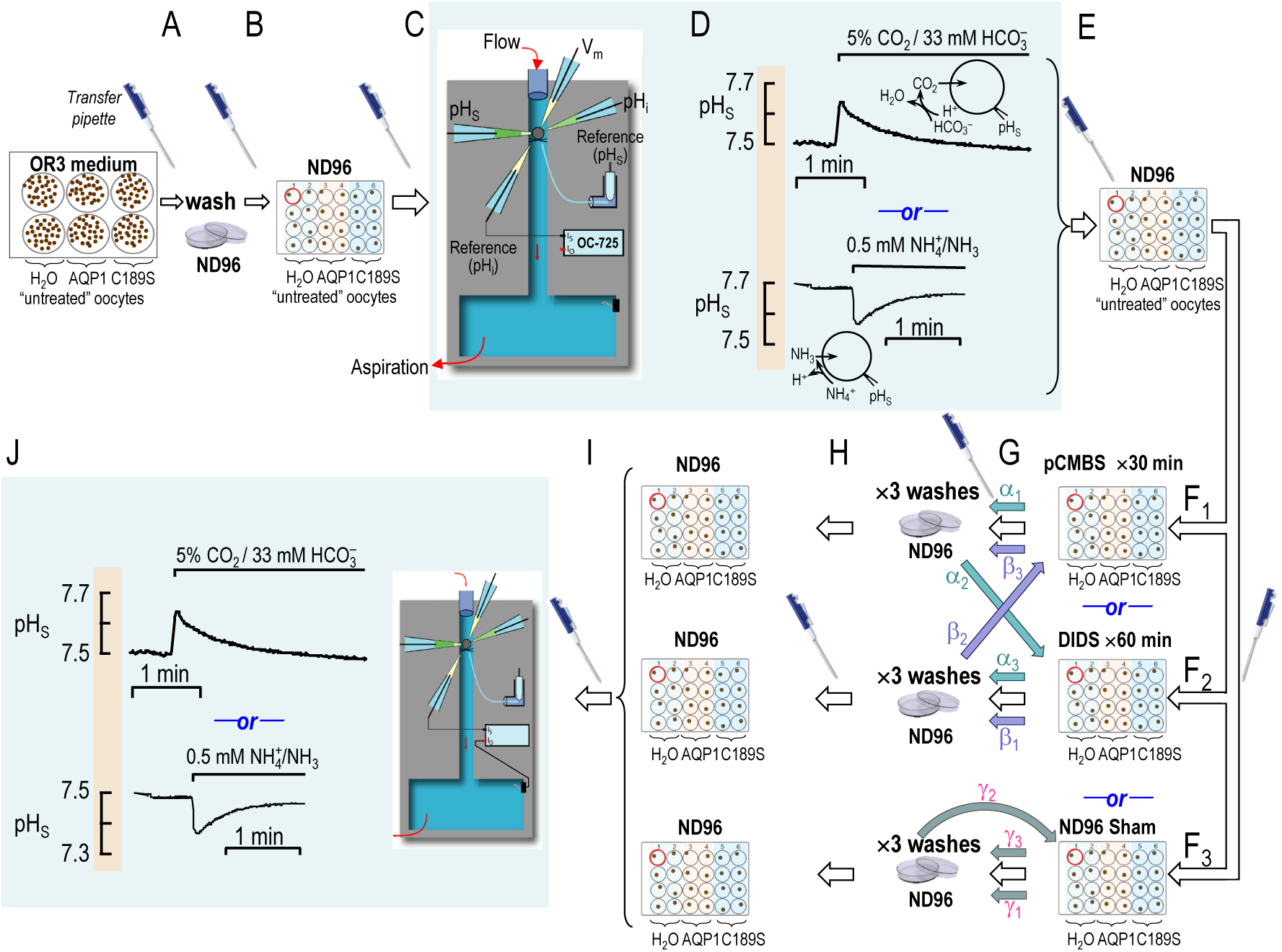
Flow chart showing experimental protocols. The letters “A”, “B”, “C” and so on are above broad arrows and near icons of a pipetter that identify the transfer of an oocyte (using a transfer pipette) from one step in our protocol to another. “D” and “J” are the initial and final electrophysiological recordings, respectively. The oocytes may be controls (injected with H_2_O) or those expressing either hAQP1-WT (AQP1) or hAQP1-C189S (C189S). In transfer “F”, we move the oocyte to a solution containing pCMBS (F_1_), DIDS pCMBS (F_2_), or no drug (F_3_). As indicated by the white arrows, we can directly transfer from F_1_ (or F_2_ or F_3_) to G to H. Alternatively, at transfer “F”, we can execute subsidiary protocols (α/teal arrows, β/lavender arrows, γ/gray arrows) in which we return oocytes from washes in ND96 back to a solution containing an inhibitor (i.e., pCMBS or DIDS) or ND96 (i.e., sham drug exposure). Thus, an oocyte might undergo the teal-colored pCMBS/DIDS sequence (α_1_α_2→_α_3_), the lavender-colored DIDS/pCMBS sequence (β_1_β_2→_β_3_), or the gray-colored Sham/Sham sequence (γ_1_γ_2→_γ_3_). After transfer H we perform the final electrophysiology recordings.

#### Electrophysiological assays

On the day of an experiment, we followed a 10-step protocol:

A. Use a transfer pipette to pick up an untreated oocyte from a well of a 6-well plate (each well containing OR3 medium + ∼30 oocytes, all injected with the same material) and drop the oocyte into a petri dish containing ND96 (Solution #1, Table 1). After flushing the transfer pipette tip several times with ND96 (to remove OR3).
B. Use the transfer pipette to move the washed oocyte to an identified well of a 24-well plate (1 oocyte/well) containing ND96. Repeat this procedure for several oocytes, each of which will remain in its well (1–3 hours) until we …
C. Use the transfer pipette to place the oocyte in the chamber, begin flowing ND96, and measure *V*_m_, pH_i_, and pH_S_ under basal conditions.
D. Monitor pH_S_ as we replace ND96 with either the 5% CO_2_/33 mM 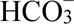 solution (Solution #3, Table 1; as in Figure 2B, Figure 3) or the 0.5 mM NH_3_/NH_4_^+^ solution (Solution #5, Table 1; as in Figure 5B, Figure 6); all oocytes in the study underwent this control assay.

**Figure 2:**
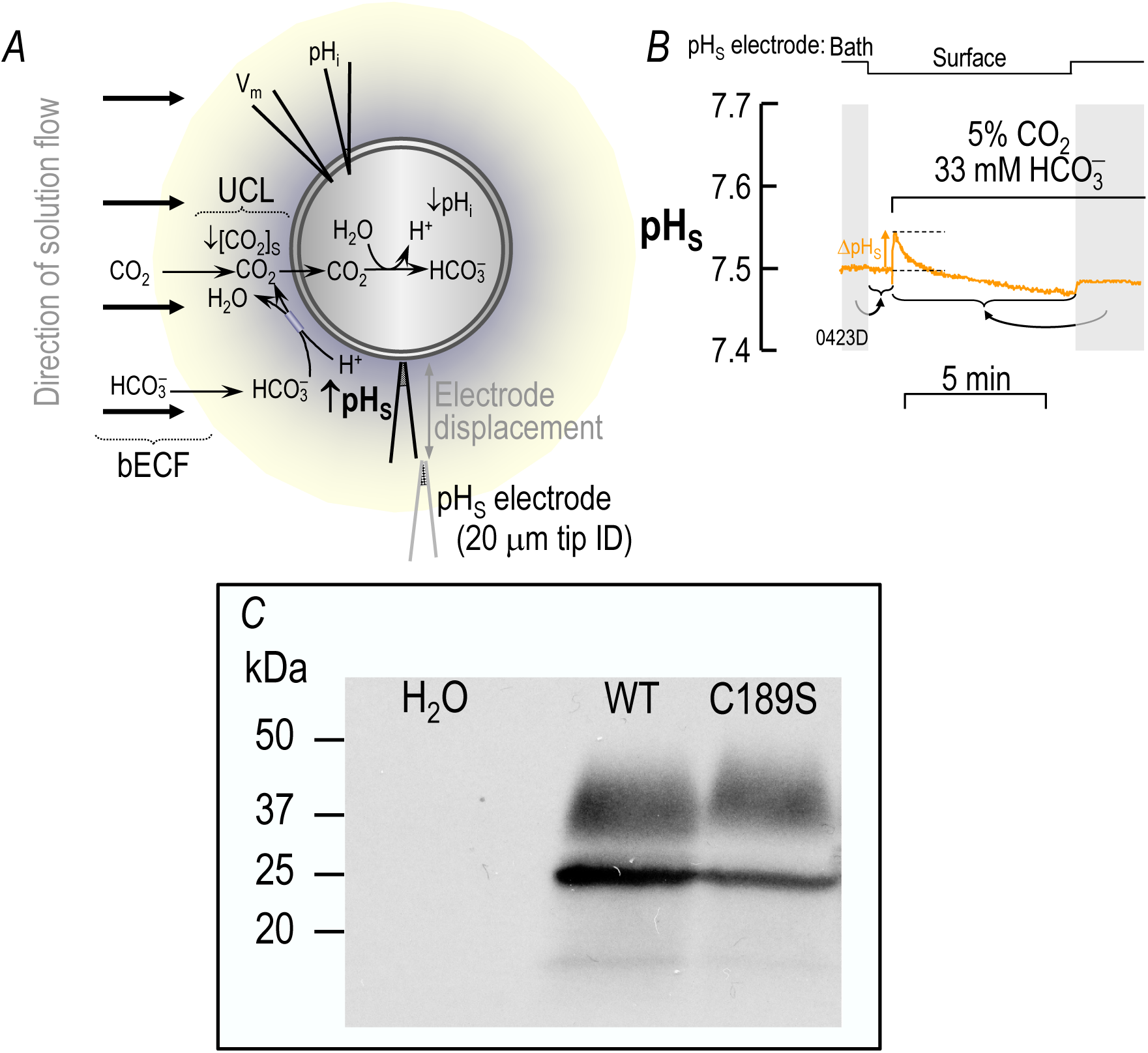
CO_2_ influx into an oocyte. *A,* schematic illustration of a CO_2_-influx experiment. Thick black arrows indicate direction of convective flow within the bulk extracellular fluid (bECF). Thinner arrows indicate solute diffusion or reactions. The maize-colored halo indicates the layer of extracellular unconvected fluid (EUF). At the outer surface of the membrane, CO_2_ influx creates a CO_2_ deficit, in part replenished by the reaction 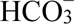 + H^+^ → CO_2_ + H_2_O (which raises pH_S_) and in part replenished by diffusion from the bECF (an isohydric process). The double-headed arrow indicates displacement of the pH_S_ electrode from the cell surface to the bECF for recalibration. *B,* sample surface-pH (pH_S_ record). The upward arrow indicates the maximal change in pH_S_ (ΔpH_S_) during the application of CO_2_/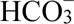. During the two periods indicated by the vertical gray bands, the tip of the pH_S_ electrode was displaced ∼300 μm away from the cell surface for recalibration in the bulk extracellular fluid (i.e., pH = 7.50). The curved gray arrow points to horizontal braces that indicate the portions of the experiment to which each calibration pertains. The filename for this representative trace, “0423D”—that we reuse in Figure 3A—is annotated at the bottom left corner of the panel. *C,* western blot, probed with an anti-AQP1 antibody, showing relative expression of AQP1-WT vs. AQP1-C189S in one of 3 membrane preparation of oocytes. ID, inner diameter; UCL, unconvected layer.

**Figure 3:**
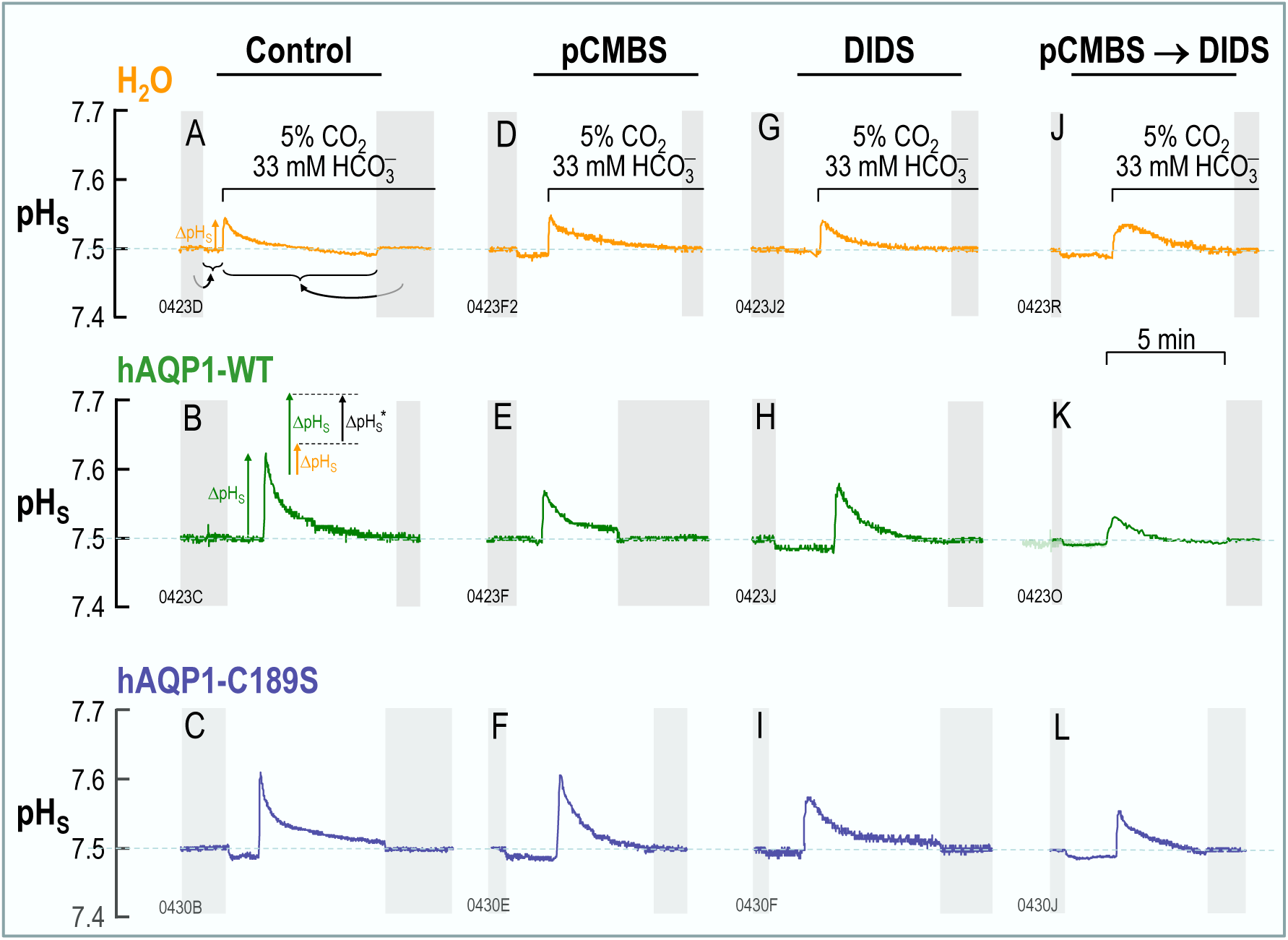
Surface-pH transients triggered by a CO_2_/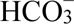 exposure and subsequent CO_2_ influx. The figure shows representative pH_S_ records from oocytes injected with H_2_O (top row) or cRNA encoding hAQP1-WT (middle row) or its C189S mutant (bottom row), and later exposed first to the ND96 solution and then, at the indicated times, to a solution containing CO_2_/HCO_3_^−^. All oocytes underwent the “Control” protocol, followed by one or two of the three indicated F_1_/F_2_/F_3_ protocols in Figure 1, although some oocytes did not survive beyond the control protocol. As necessary, we pre-incubated oocytes in 1 mM pCMBS for 30 minutes, or in 100 µM DIDS for 1 hour. Neither drug was present in the bulk solution at the time of the assays. The two gray bars in each panel indicate when we withdrew the pH-electrode tip from the oocyte surface to the bulk extracellular fluid (pH 7.50) for recalibration. In panel *A*, the curved gray arrows points and horizontal braces have the same meanings as in Figure 2B, and the upward gold arrow indicates the control ΔpH_S_. In panel *B*, the upward green arrow indicates the ΔpH_S_ in an oocyte expressing AQP1. The inset shows that the difference in heights of the green arrow and the gold arrow (its day-matched control) is the channel-dependent ΔpH_S_ (ΔpH_S_*).

**Figure 4:**
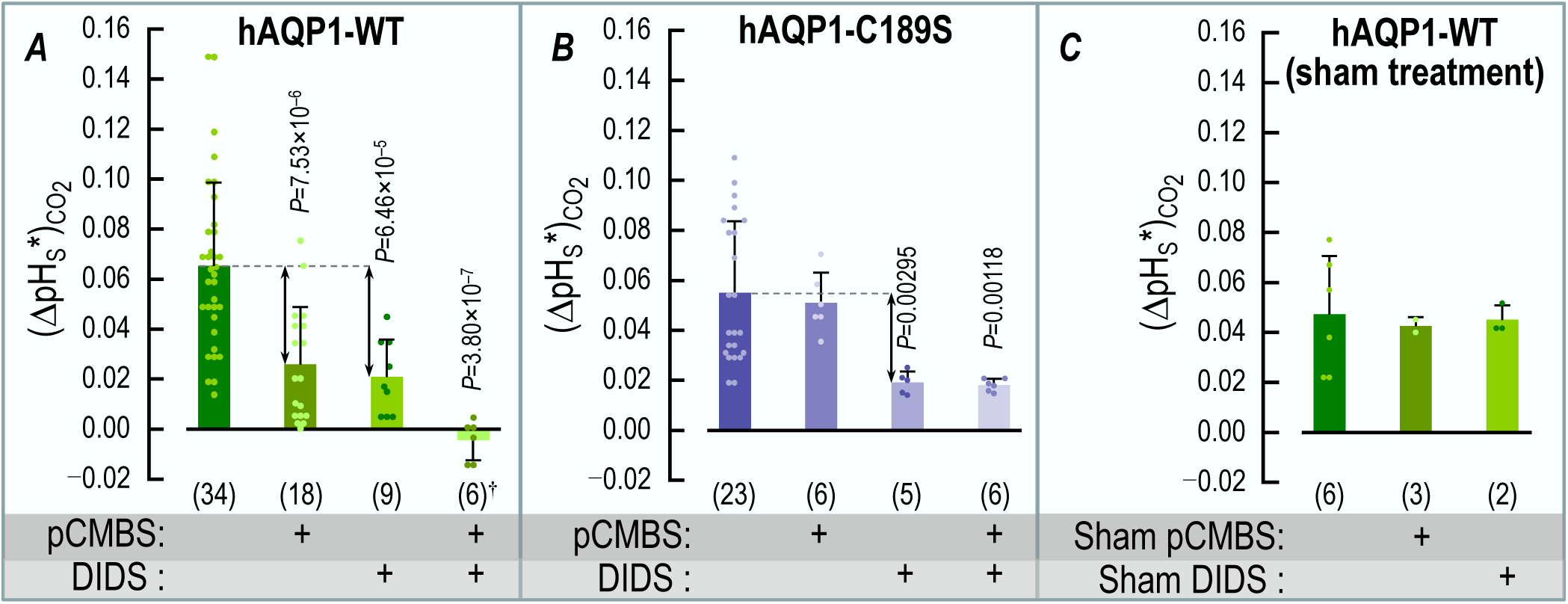
Summary of channel-specific data in assays for CO_2_ influx. *A,* oocytes expressing hAQP1-WT. pCMBS and DIDS each reduces (ΔpH_S_*)_CO2_ by somewhat more than half, and the two drugs together reduce it to virtually zero. For the rightmost bar, the dagger symbol (†) adjacent to the replicates (i.e., n = 6) indicates that the (ΔpH_S_*)_CO2_ for the pCMBS+DIDS condition is not significantly different from zero (*P*=0.254, one-sample t-test). Of the 6 pCMBS+DIDS oocytes, 2 we treated first with pCMBS, and then DIDS; the other 4 were in the opposite order. *B,* oocytes expressing the C189S mutant of hAQP1. DIDS reduces (ΔpH_S_*)_CO2_ by about half, but pCMBS is without effect, ±DIDS. Of the 6 pCMBS+DIDS oocytes (far-right bar), 4 we treated first with pCMBS, and then DIDS; the other 2 were in the opposite order. *C,* oocytes expressing AQP1 but undergoing only sham drug exposures. These sham experiments show that the long protocols did not have a substantial effect on (ΔpH_S_*)_CO2_. The pCMBS shams (30 min) and DIDS shams (60 min) differed only in the ND96 incubation time in Figure 1/F_3_. The data come from experiments like those in Figure 3, in which we expose oocytes to 5% CO_2_/33 mM 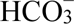 (for protocol, see Methods and Figure 1). From each ΔpH_S_ of a channel-expressing oocyte (e.g., Figure 3B, *E*, *H*, *K* or *C*, *F*, *I*, *L*), we subtract the mean, day-matched ΔpH_S_ for the corresponding H_2_O-injected oocytes (e.g., Figure 3A, *D*, *G*, and *J*) to calculate the channel-dependent ΔpH_S_ for CO_2_, that is, (ΔpH_S_*)_CO2_. The (ΔpH_S_*)_CO2_ values from individual oocytes are plotted as dots over the green-shaded bars in panels *A* and *C*, and purple-shaded bars in panel *B*. At the base of each bar in parentheses is the numbers of oocytes (n), which come from a minimum of 5 batches of oocytes (i.e., different frogs; N ≥ 5). Error bars represent S.D. In the horizontal gray bands at the bottom of each panel, “+” indicates a pre-incubation with pCMBS or DIDS, or a sham exposure. *P*-values denote statistically significant differences from the no-drug condition, and are results of one-way ANOVAs among all groups, followed by Holm-Bonferroni post hoc means comparisons (see Statistics in Methods). For clarity, we display only *P*-values that indicate statistical significance; we show the *P*-values for all comparisons in Statistics Table 4*a*, Statistics Table 4*b* and Statistics Table 4*C*.

**Figure 5:**
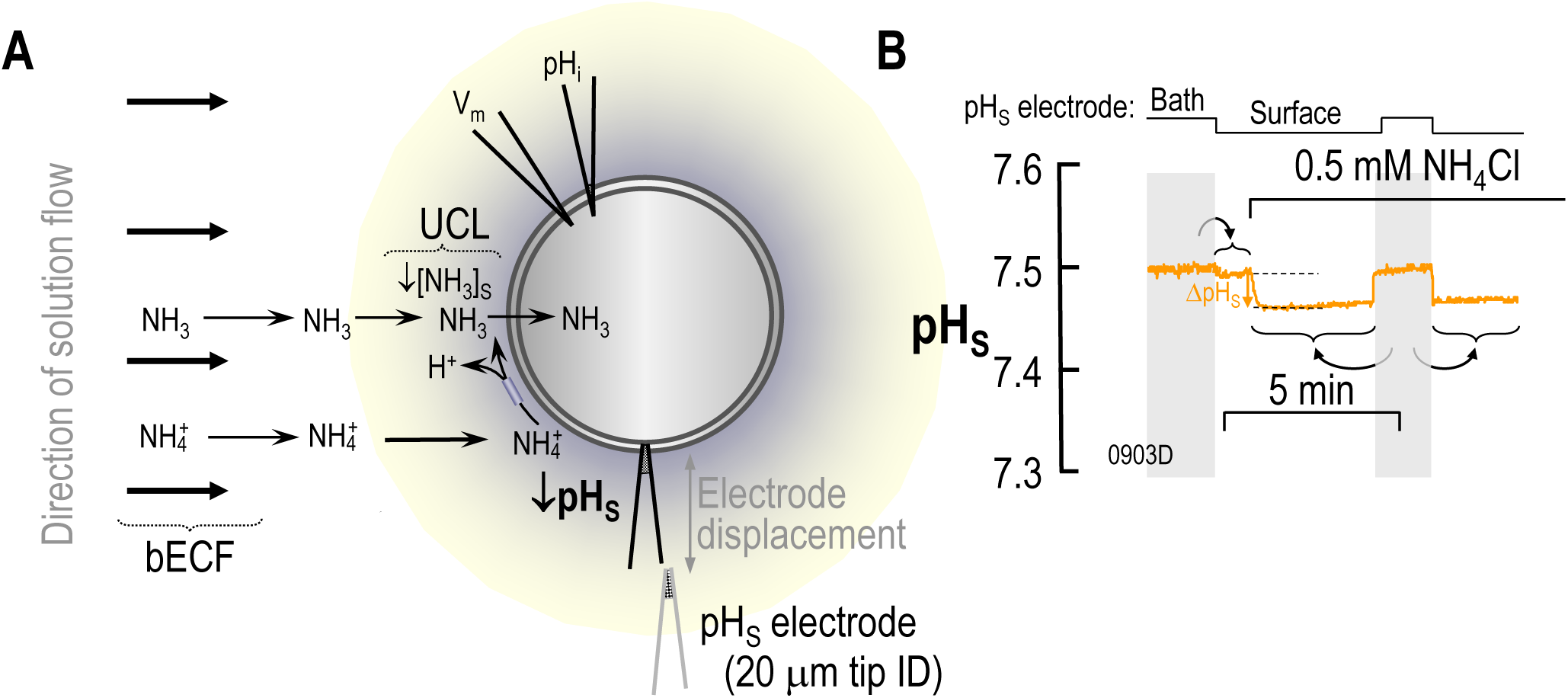
NH_3_ influx into an oocyte. *A,* schematic illustration of a NH_3_-influx experiment. Analogous to Figure 2, the thick black arrows indicate the direction of convective flow within the bulk extracellular fluid (bECF). Thinner arrows indicate solute diffusion or reactions. The maize-colored halo indicates the layer of extracellular unconvected fluid (EUF). At the outer surface of the membrane, NH_3_ influx creates a NH_3_ deficit, in part replenished by the reaction NH^+^ → NH_3_ + H^+^ (which decreases pH_S_) and in part replenished by diffusion from the bECF (an isohydric process). The double-headed arrow indicates movement of the pH_S_ electrode from the bECF to the cell surface for recalibration. *B,* Example of a surface-pH (pH_S_) record. The downward arrow indicates the maximal change in pH_S_ (ΔpH_S_) during the application of NH_3_/NH^+^. During the two periods indicated by the vertical gray bands, the tip of the pH_S_ electrode was ∼300 μm away from the cell surface for calibration in the bulk extracellular fluid (i.e., pH = 7.50). The curved gray arrows point to horizontal braces that indicate the portions of the experiment to which each calibration pertains. ID, inner diameter. The filename for this representative trace, “0903D”—that we reuse in Figure 6A—is annotated at the bottom left corner of the panel.

**Figure 6:**
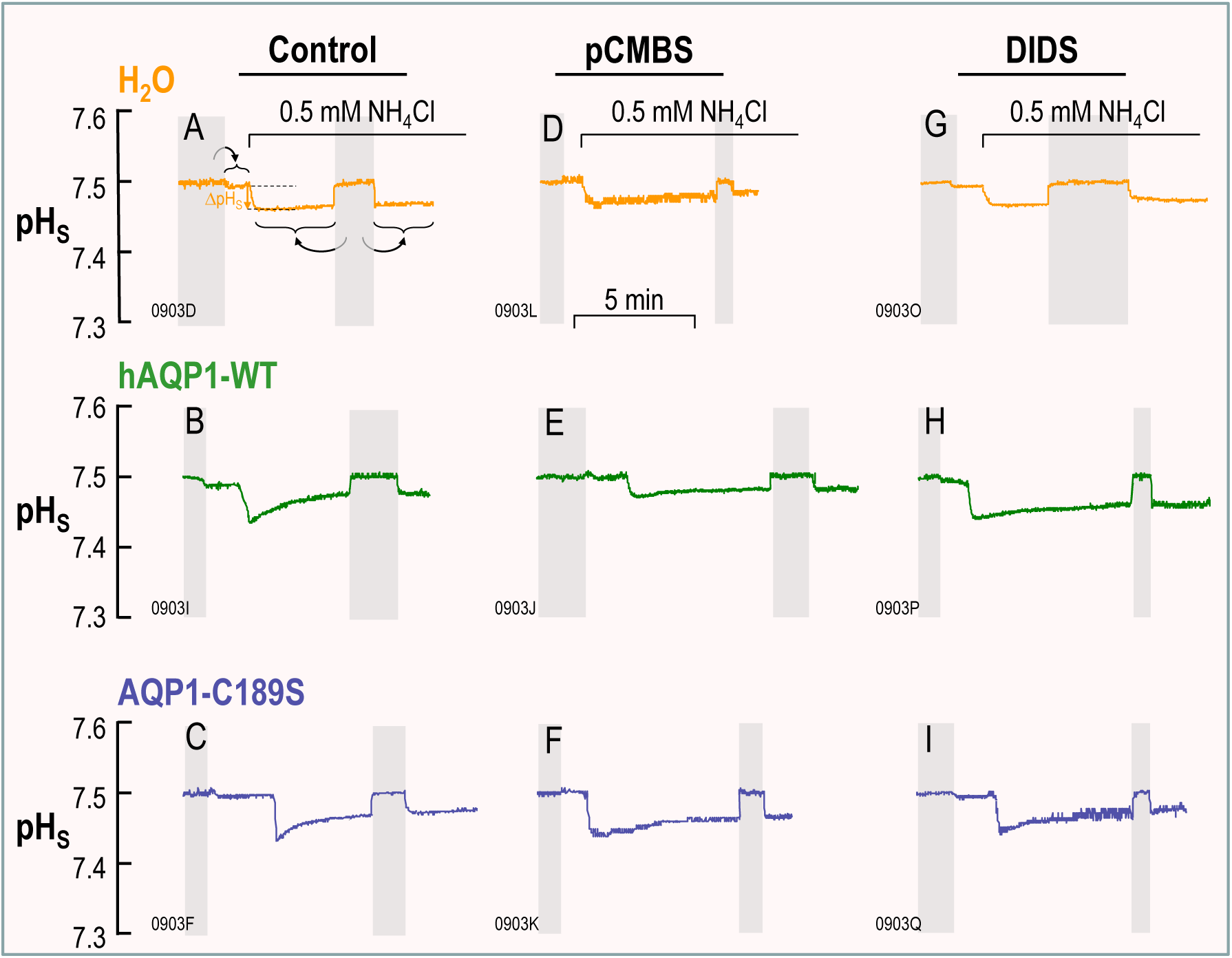
Surface-pH transients triggered by NH_3_/NH^+^ exposure and the subsequent NH_3_ influx. The figure shows representative pH_S_ records from oocytes injected with H_2_O (top row) or cRNA encoding hAQP1-WT (middle row) or its C189S mutant (bottom row), and later exposed first to the ND96 solution and then, at the indicated times, to a solution containing NH_3_/NH_4_^+^. All oocytes underwent the “Control” protocol, followed by the F_1_ or F_2_ protocol in Figure 1, although some oocytes did not survive beyond the control protocol (see Methods). As necessary, we pre-incubated oocytes in 1 mM pCMBS for 30 min, or in 100 µM DIDS for 1 h. Neither drug was present in the bulk solution at the time of the assays. The gray bars indicate when we withdrew the pH-electrode tip from the oocyte surface to the bulk extracellular solution (pH 7.50) for recalibration. In each panel the filename the representative trace, is annotated at the bottom left corner of the panel. “0903D” in panel *A* is a reproduction of the recording in Figure 5B.
E. Immediately upon completion of the pH_S_ assay, use a transfer pipette to return the oocyte from the chamber to its original well and incubate the oocyte at ∼22°C in ND96. We serially process all oocytes from a 24-well plate up to this stage (i.e., steps ‘C’ through ‘E’) and then pause. Thus, this second period of incubation in ND96 (step ‘E’) ranges from ∼30 min to ∼3 h.
F. Use a transfer pipette to move each oocyte in the 24-well ND96 plate to a corresponding well in one of three new 24-well plates. Here, the wells contain one of the following three solutions: (F_1_) 1 mM pCMBS dissolved in ND96 (Solution #7, Table 1) 30 min incubation; see Figure 3DEF, Figure 6DEF, Figure 7, Figure 8, or Figure 9CD); (F_2_) 100 µM DIDS dissolved in ND96 (Solution #8, Table 1; 60 min incubation; see Figure 3GHI or Figure 6GHI); or (F_3_) ND96 without an added drug (“sham”).

**Figure 7:**
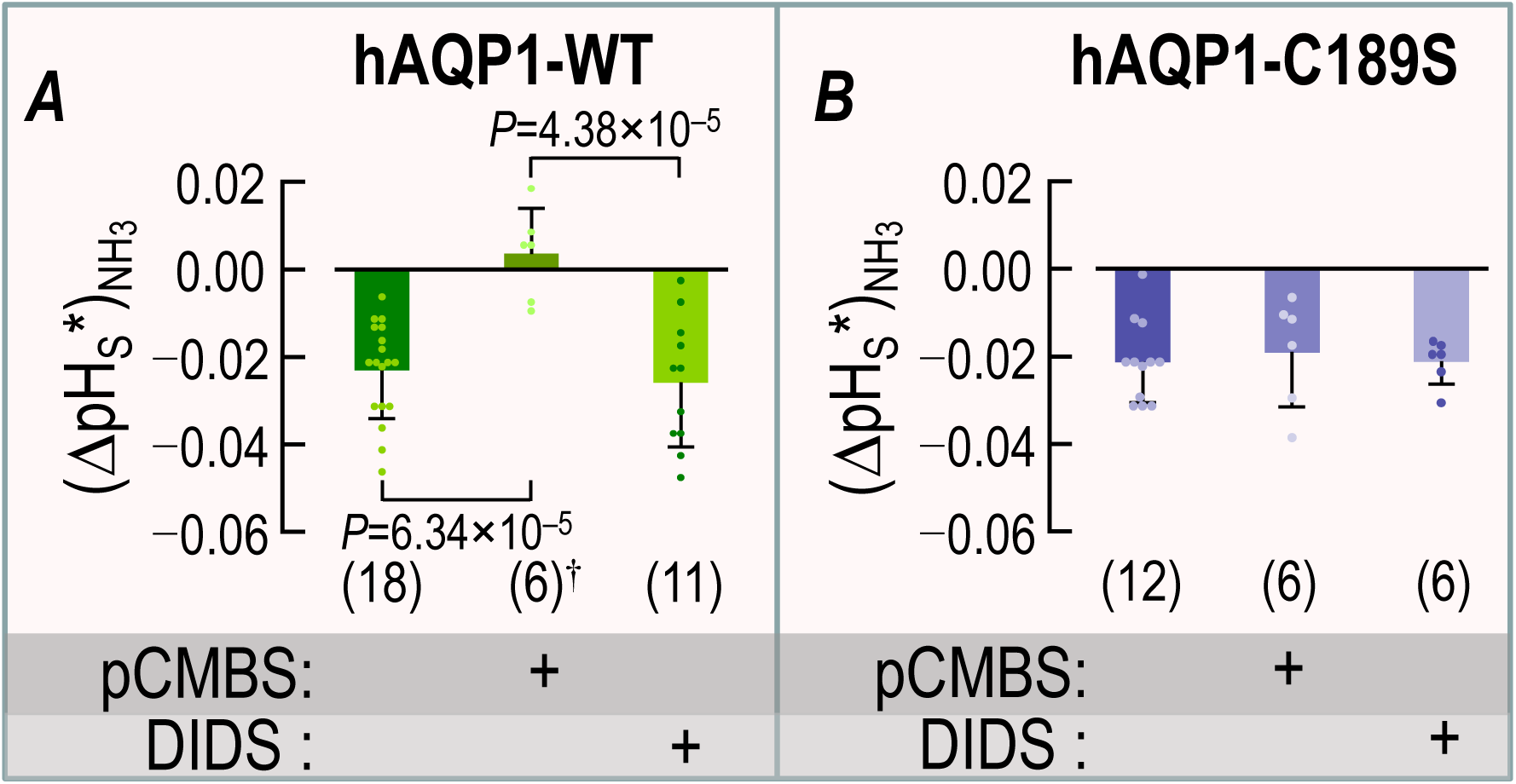
Summary of channel-specific data in assays for NH_3_ influx. *A*, oocytes expressing hAQP1-WT. pCMBS reduces (ΔpH_S_*)_NH3_ to virtually zero, whereas DIDS has no effect. For the middle bar, the dagger symbol (†) adjacent to the replicates (i.e., n = 6) indicates that the (ΔpH_S_*)_NH3_ for the pCMBS condition is not significantly different from zero (*P*=0.450, one-sample t-test). *B*, oocytes expressing the C189S mutant of hAQP1. Neither drug affects (ΔpH_S_*)_NH3_. The data come from experiments like those in Figure 6, in which we exposed oocytes to 0.5 mM NH_4_Cl (for protocol, see Methods and Figure 1). From each ΔpH_S_ of a channel-expressing oocyte (Figure 6B, *E*, *H* or *C, F, I*), we subtracted the corresponding mean, day-matched ΔpH_S_ for H_2_O-injected oocytes (Figure 6A*, D, G*) to calculate the channel-dependent ΔpH_S_ for NH_3_, that is, (ΔpH_S_*)_NH3_. The (ΔpH_S_*)_NH3_ values from individual oocytes are plotted over the green-shaded bars in panel *A*, and purple-shaded bars in panel *B*. At the base of each bar in parentheses is the number of oocytes (n), which come from a minimum of 5 batches of oocytes (i.e., different frogs; N). Error bars represent S.D. In the horizontal gray bands at the bottom of each panel, “+” indicates a pre-incubation with pCMBS or DIDS. *P*-values denote statistically significant differences from the no-drug condition, and are the results of one-way ANOVAs among all groups, followed by Holm-Bonferroni post-hoc means comparisons (see Statistics in Methods). For clarity, we display only *P*-values that indicate statistical significance; we show *P*-values for all comparisons in Statistics Table 7*a* and Statistics Table 7*b*.

**Figure 8:**
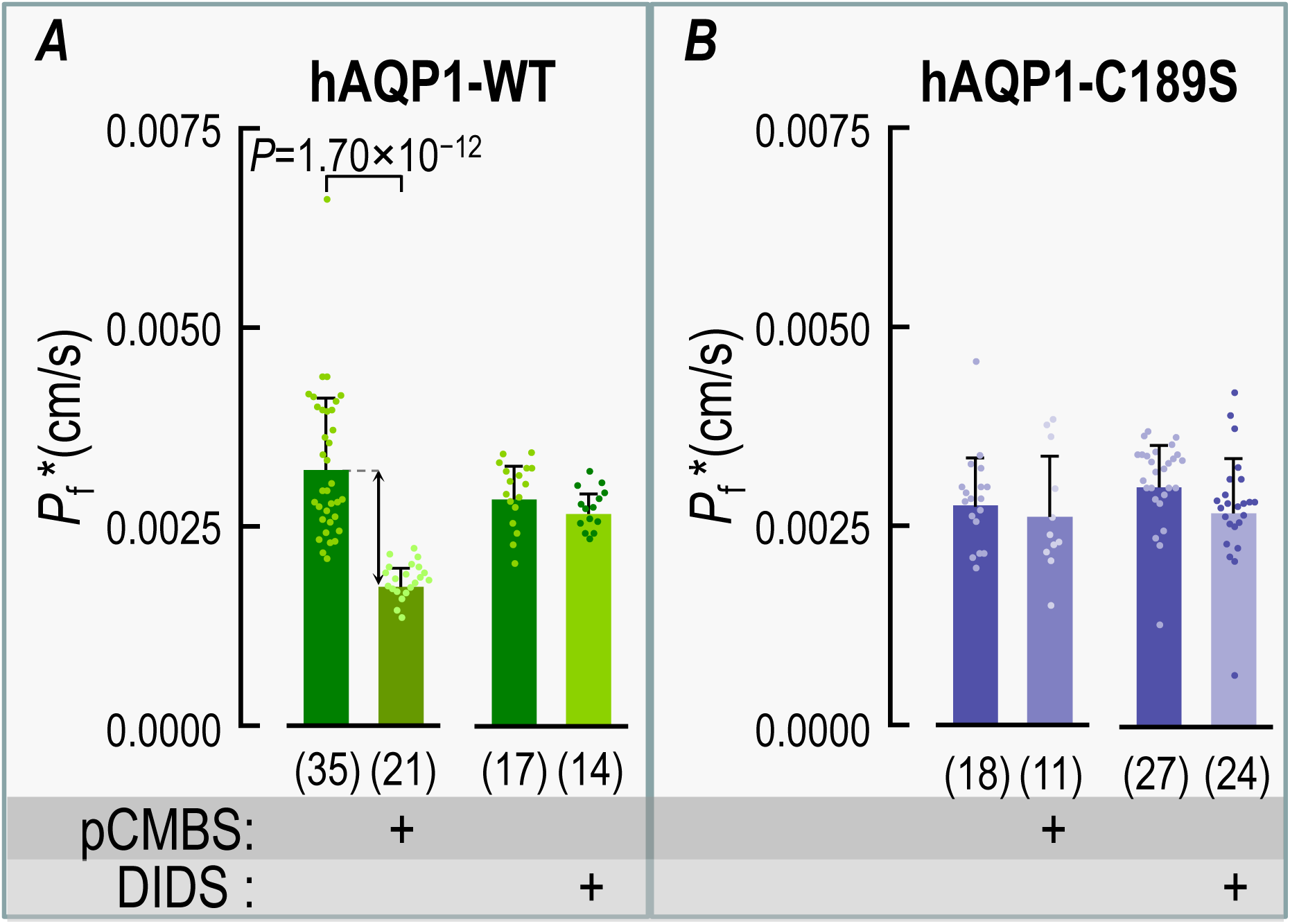
Summary of channel-specific data in assays for osmotic water permeability. *A*, oocytes expressing hAQP1-WT. pCMBS but not DIDS reduces *P*_f_. *B*, oocytes expressing the C189S mutant of hAQP1. Neither drug reduces *P*_f_. From each *P*_f_ from a channel-expressing oocyte, we subtracted the corresponding mean, day-matched *P*_f_ for H_2_O-injected oocytes to calculate the channel-dependent *P*_f_, that is, *P*_f_*. The *P*_f_* values from individual oocytes are plotted over the green-shaded bars in panel *A*, and purple-shaded bars in panel *B*. At the base of each bar in parentheses is the number of oocytes (n), which come from a minimum of 5 batches of oocytes (i.e., different frogs; N). Error bars represent S.D. In the horizontal gray bands at the bottom of each panel, “+” indicates a pre-incubation with pCMBS or DIDS. *P*-values denote statistically significant differences from the no-drug condition, and are the results of a one-way ANOVAs among all groups, followed by Holm-Bonferroni post-hoc means comparisons (see Statistics in Methods). For clarity, we display only *P*-values that indicate statistical significance; we show *P*-values for all comparisons in Statistics Table 8.
G. Wash the oocyte. Except in a few cases (see indented paragraph below), we wash the oocyte ×3 in a petri dish containing ordinary ND96, as follows. We use a transfer pipette to pick up an oocyte from a 24-plate (step ‘F’) and release the oocyte into a petri dish containing ND96, quickly pick up and released the oocyte into this wash solution twice more (to remove excess drug, if present). At this point, most oocytes progress immediately through step ‘H’ (below). However, in a minority of experiments (see Figure 3JKL), we serially treated oocytes with both pCMBS and DIDS. Here, oocytes previously treated with pCMBS (step ‘F_1_’) we exposed to DIDS, as indicated by steps α_1_α_2_α_3_. Conversely, oocytes previously treated with DIDS (step ‘F_2_’) we exposed to pCMBS, as indicated by steps β_1_β_2_β_3_. In one experiment without any drugs, we also followed the steps γ_1_γ_2_γ_3_ (“double sham”). Not illustrated in Figure 1 is a protocol variant that we employ in a few cases (i.e., Figure 9AB*/right bars*) in which we introduce an additional step between ‘F_2_’ and ‘G’. Here, after treating an oocyte with DIDS (step ‘F_2_’), we scavenge unreacted DIDS with albumin as follows. After using a transfer pipette to pick up the oocyte from the DIDS-containing ND96 solution, we release the oocyte into a petri dish containing 0.2% albumin in otherwise DIDS-free ND96 (Solution #9, Table 1), and allow the oocyte to remain in the albumin solution for 20 min. We then wash off the albumin in step ‘G’ by transferring the oocyte to a petri dish containing ordinary ND96, and pick up and release the oocyte ×3 using a transfer pipette.
H. Use a transfer pipette to move the oocyte from the petri dish (step ‘G’) to an identified well in one of three new 24-well plates, all containing ND96, in preparation for the next assay.
I. Transfer the oocyte back to the chamber and (as in step ‘C’), and again measure the new basal *V*_m_, pH_i_, and pH_S_.
J. Monitor (as in step ‘D’), for the second time, the pH_S_ transient as we replace ND96 with the CO_2_/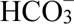 (or NH_3_/NH^+^) solution.

**Figure 9:**
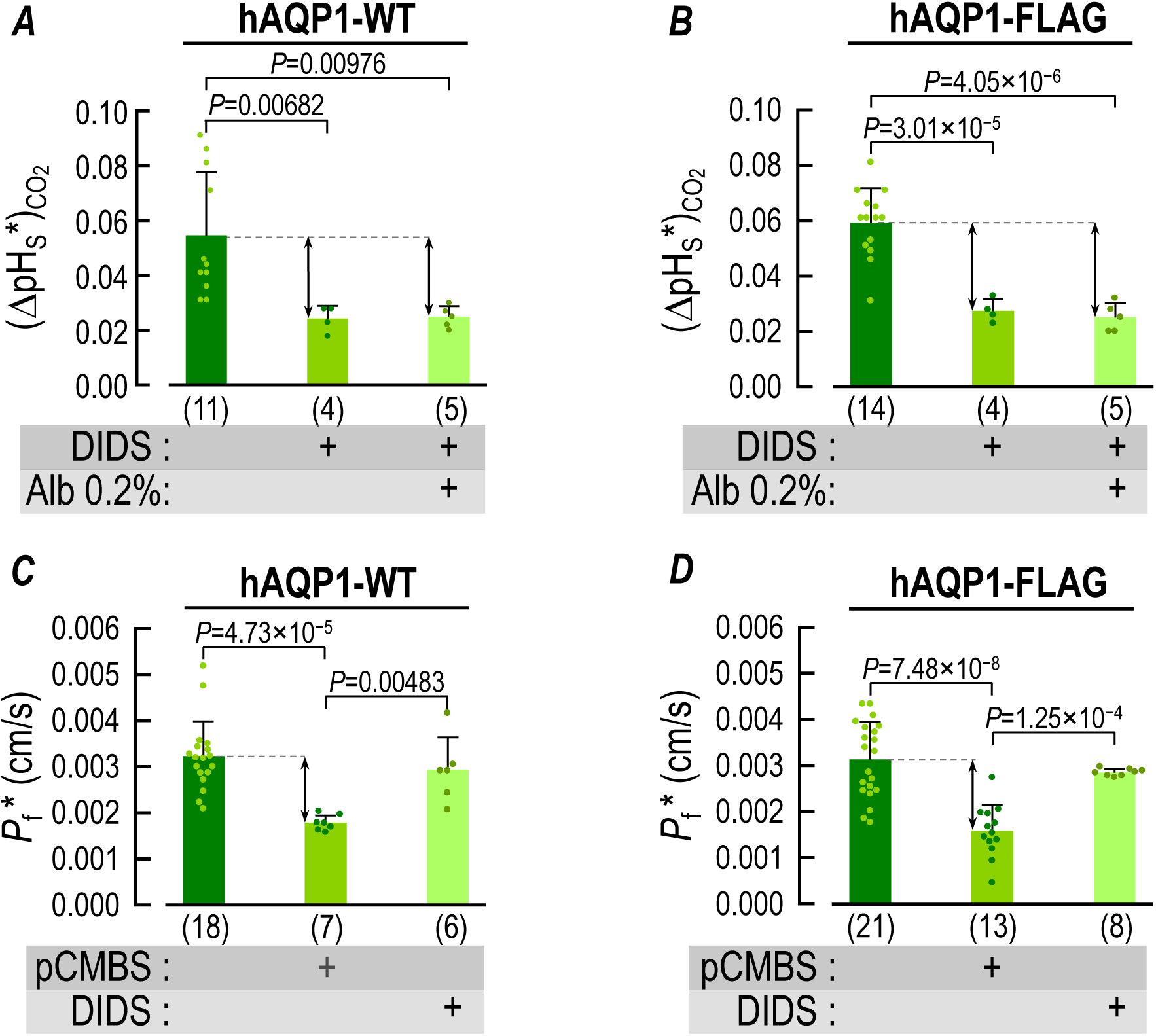
Summary of channel-specific data for CO_2_ and H_2_O influx for WT vs. FLAG-tagged hAQP1. *A,* oocytes expressing hAQP1-WT: effect of albumin washes on the DIDS sensitivity of (ΔpH_S_*)_CO2_. DIDS reduces (ΔpH_S_*)_CO2_ by about half, regardless of the albumin wash. *B,* oocytes expressing N-terminally FLAG-tagged hAQP1: effect of albumin washes on the DIDS sensitivity of (ΔpH_S_*)_CO2_. Also in the FLAG-tagged construct, DIDS reduces (ΔpH_S_*)_CO2_ by about half, regardless of the albumin wash. *C,* oocytes expressing hAQP1-WT: effect of albumin washes on the DIDS sensitivity of *P*_f_*. These experiments are a matched control for the study in panel *D*. As in the comparable experiments summarized in Figure 8A, pCMBS but not DIDS reduces *P*_f_. *D,* oocytes expressing FLAG-tagged hAQP1: effect of albumin washes on the DIDS sensitivity of *P*_f_*. As in panel *C* (no FLAG tag), pCMBS but not DIDS reduces *P*_f_ in oocytes expressing FLAG-tagged hAQP1. For panels *A* and *B*, the control pH_S_ data (leftmost dark-green bars) come from experiments like those in Figure 3, whereas for panels *C* and *D*, the control *P*_f_ data come from experiments like those in Figure 8. The (ΔpH_S_*)_CO2_ or *P*_f_* values from individual oocytes are plotted as dots over the green-shaded bars in panels *A* through *D*. At the base of each bar in parentheses is the numbers of oocytes (n), which come from a minimum of 5 batches of oocytes (i.e., different frogs; N ≥ 5). Error bars represent S.D. In the horizontal gray bands at the bottom of each panel, “+” indicates a pre-incubation with pCMBS or DIDS, or the presence of 0.2% bovine serum albumin (Alb 0.2%) *P*-values denote statistically significant differences from the no-drug condition, and are results of one-way ANOVAs among all groups, followed by Holm-Bonferroni means comparisons (see Statistics in Methods). For clarity, we display only the *P*-values that indicate statistical significance; we show *P*-values for all comparisons in Statistics Table 9.

Thus, oocytes in the top row of Figure 3 (i.e., Figure 3ADGJ) went through one of the following two sequences: Panels *A*→*D* or *A*→*J*, which requires 12–18 h for an entire group of 24 oocytes in a day’s work. Not all oocytes survived the entire protocol (as judged by the color/integrity of the animal pole).

#### *P*_f_ assays

On the same day that we performed our pH_S_ assays, we also performed *P*_f_ assays on separate cells from the same oocyte batch (i.e., oocytes prepared from the same ovary). On the same day as pH_S_ experiments, we: (1) Use a transfer pipette to remove 6 untreated oocytes of each experimental group—hAQP1, C189S mutant, or hAQP1 FLAG-tagged—from an identified 6-well plate containing ND96 (i.e., a total of 18). (2) Place the oocytes—as quickly as possible—in a Petri dish containing hypotonic ND96 (Solution #6, Table 1). (3) Use a video camera (described above) to monitor cell swelling; oocytes of each group underwent this control assay. (4) Immediately remove the oocytes from the hypotonic solution to identified wells in another Petri dish containing standard ND96 solution to wash the oocytes ×3 at ∼22°C (described above under electrophysiological assays). (5) Incubate at ∼22°C either for (5a) 30 min in ND96 containing 1 mM pCMBS (Solution #7, Table 1), or (5b) 1 hour in ND96 containing 100 µM DIDS (Solution #8, Table 1). (6) Wash the oocytes (as described under electrophysiological assays, point ‘<u>5a</u>’). (7) Transfer the oocytes to a Petri dish containing hypotonic, drug-free ND96 as in ‘2’ above. (8) Monitor swelling as in ‘3’ above.

### Generation of the DIDS antibody

#### Generation of DIDS-fusion protein

We used two rabbits to produce polyclonal antibodies against DIDS, as described by Garcia & Lodish (1989). Briefly, we reacted 2.4 mM DIDS (catalog # D3514, Sigma-Aldrich), with 5 mg keyhole limpet hemocyanin (KLH;, catalog # H7017, Sigma-Aldrich) in 500 μL of phosphate buffer saline (PBS; i.e., 150 mM NaCl, 10 mM phosphate buffer, pH 7.0) in the dark at 37°C × 40 min. Afterwards, we centrifuged the sample × 30 s in an Eppendorf benchtop centrifuge (catalog # 5415C, TFS) to remove large insoluble aggregates After pooling yellow supernatants from several centrifuge tubes, we dialyzed against PBS at 4°C overnight in a slide-a-lyzer dialysis cassette (TFS). The following day, we removed the KLH-DIDS sample from the cassette; quantified the DIDS by absorbance spectroscopy at 340 nm, assuming a molar extinction coefficient (ε) of 54,000 M^−1^ cm^−1^; aliquoted the dialyzed KLH-DIDS solution; and immediately used it for immunization.

For both rabbits, we separately processed each of three blood samples (see above)—2 lots of ∼30–50 mL plus the terminal sample. We centrifuged the blood for 10 min at 1000 ×g using an Eppendorf bench top centrifuge to isolate the serum, and then evaluated antigenicity on western blots of DIDS-labeled AQP1. We purified the antibodies (mainly IgG) from the remainder of the serum using a Protein A column (Catalog # 20356, TFS), following the manufacturer’s instructions; estimated [IgG] by measuring absorbance at 280 nm (assuming ε = 210,000 M^-1^cm^-1^) using a NanoDrop 2000c spectrophotometer (TFS); separated the material into aliquots corresponding to 1 mg/mL of anti-DIDS antibody; and stored the aliquots at either –20°C for short-term storage or –80°C for long-term storage.

### Biochemical experiments

#### Isolation of oocyte membranes

We used a previously described method (Leduc-Nadeau *et al*., 2007), except that, after disrupting oocytes and centrifuging, we removed the supernatant and added fresh buffer to the pellet, repeating the procedure until the supernatant was clear (i.e., void of yolk).

#### Purification of oocyte membrane proteins

We resuspended isolated oocyte membranes in Tris-Buffered Saline (TBS, 50 mM Tris HCl, pH 7.4, 150 mM NaCl) + 2% n-dodecyl-β-D-maltopyranoside (DDM; Sol-Grade, catalogue #D310S, Anatrace, Maumee, OH), incubated at 4°C × 2h, centrifuged at 16,000 × g at 4°C × 30 min, and then collected the supernatant containing the solubilized membrane proteins and diluted it so that [DDM] was ≤ 0.5%.

#### Overexpression of AQP1-FLAG in *Pichia pastoris*

We ligated N-terminally FLAG-tagged hAQP1 (M<u>DYKDDDDKASEFKKKL…</u>; FLAG sequence underscored, hAQP1 sequence double underscored) into the pPICZ-A vector (Invitrogen, Carlsbad, CA), induced protein expression with methanol × 72 h at 26 °C, harvested the yeast by centrifugation, and obtained spheroplasts using zymolyase treatment (60 U/mL in 1 M sorbitol, 1 mM EDTA, 10 mM citrate buffer, pH 5.8) as previously described (Gustin *et al*., 1998). We then equally divided the spheroplasted cells into two groups (i.e., no DIDS and added 100 µM DIDS), incubated × 1 h at ∼22 °C, and then solubilized overnight at 4°C in TBS + 2% DM, with cOmplete™ Protease Inhibitor Cocktail (catalog #11697498001, Roche, Mannheim, Germany). Following membrane solubilization, we clarified the material by low speed centrifugation, mixed with anti-FLAG resin (catalog #A2220, Sigma-Aldrich), and eluted proteins from a column using TBS + 0.1% DM + 500 µg/mL FLAG peptide (catalog #F4799, Sigma-Aldrich) at pH 7.4. In some cases, we subjected FLAG-purified AQP1 to size-exclusion chromatography using a Sephadex 75 column.

#### Western blotting of membrane proteins from oocytes or *Pichia pastoris*

We separated proteins extracted from oocyte membranes by SDS-PAG using Tris-Glycine 4–20% gels (BioRad Laboratories, Hercules, CA) or proteins from extracted from *Pichia* membranes using 12% Tris-Glycine gels; transferred the proteins to polyvinylidene difluoride (PVDF) membranes using an iBlot 7-Minute Blotting System (TFS); and subsequently blocked the cross-linked proteins with Tris-buffered saline with tween (TBST), comprising 25 mM Tris at pH 7.4, 150 mM NaCl, 0.05% Tween-20 (catalog # P9416, Sigma-Aldrich) + 5% milk powder × 1 h.

We probed blots from oocyte samples with a polyclonal anti-AQP1 (catalog # AQP11-A, Alpha Diagnostics, San Antonio, USA), and blots from *P. pastoris* samples with monoclonal anti-FLAG (catalog # F3165-2MG, Sigma-Aldrich) or polyclonal anti-DIDS (made in-house, see above), applying all primary antibodies at 1:1000 dilution from their stocks in TBST + 5% milk, at 4°C overnight. The next day, we washed the blot ×5 with TBST × 10 min. We detected the primary polyclonal antibodies (i.e., anti-AQP1 and anti-DIDS) with an HRP-conjugated goat anti-rabbit secondary antibody (catalog # 1706515, BioRad Laboratories), and detected the primary monoclonal antibody (i.e., anti-FLAG), an HRP-conjugated goat anti-mouse secondary antibody (catalog # STAR207P, BioRad Laboratories). We applied both secondary antibodies at 1:5000 dilution from their stocks in TBST × 1 h, washed the blot ×5 with TBST × 10 min, developed the immunoblots using ECL Prime Western Blotting Detection Reagent (catalog # 12316992, GE Healthcare Amersham, Piscataway, NJ), acquired images with a Typhoon Trio gel-documentation system (GE Healthcarefryan), and evaluate band density patterns using Image J software (NIH, Bethesda, MD).

### Statistics

We present data as the mean ± S.D. (standard deviation), and define N as the number of different frogs used and n as the number of replicate experiments (i.e., oocytes) for each condition in each dataset. We define *m* as the number of comparisons in each dataset. Statistical analyses are performed using Origin 2024 software. Statistical comparisons among means were performed using a one-way analysis of variance (ANOVA) among all groups, followed by Holm-Bonferroni mean comparison (Holm, 1979) to control for type I errors across multiple comparisons, establishing the familywise error rate (FWER) to α = 0.05. In brief, the unadjusted *P*-values for all *m* comparisons in each dataset from lowest to highest. In the first test, we compare the smallest unadjusted *P*-value to the first adjusted α value, α/*m*. Upon rejection of the null hypothesis, we compare the second-smallest *P*-value to the second adjusted α value, α/(*m*–1), and continue the process accordingly. If the unadjusted *P*-value is ≥ the adjusted α at any stage, the null hypothesis is accepted, rending all subsequent hypotheses in the test group null. We conducted one-sample t-tests (α=0.05) to ascertain whether the channel-dependent (ΔpH_S_*)_CO2_ was significantly different from zero when oocytes were treated with both pCMBS and DIDS. If *P* < 0.05, the (ΔpH_S_*)_CO2_ was considered significantly greater than zero. We conducted one-sample t-tests (α=0.05) to ascertain whether the channel-dependent (ΔpH_S_*)_NH3_, was significantly different from zero when oocytes were treated with pCMBS. If *P* < 0.05, the (ΔpH_S_*)_CO2_ was considered significantly greater than zero.

## Results

### Surface-pH Measurements for AQP1-WT and AQP1-C189S

#### pH_S_ measurements for CO_2_ transport

Figure 2*a* is a schematic representation of the reaction and diffusion events that take place as we switch the bath solution—that is, the bulk extracellular fluid—from ND96 to another that contains 5% CO_2_/33 mM 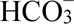 at a constant pH of 7.50. This solution change not only leads to a net influx of CO_2_ that lowers intracellular pH (Roos & Boron, 1981) but also lowers CO_2_ concentration near the outer surface of the cell membrane ([CO_2_]_S_). This decrease in [CO_2_]_S_ has two effects. First, it creates a gradient for CO_2_ diffusion from the bECF to the cell surface, which partially replenishes CO_2_ at the cell surface but by itself has no effect on pH. Second, it drives the reactions 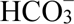 + H^+^ → H_2_ CO_3_ → CO_2_ + H_2_O at the cell surface, which also partially replenishes CO_2_. This reaction causes an alkaline pH_S_ transient, the maximal excursion of which we define as ΔpH_S_ (Endeward *et al*., 2006; Musa-Aziz *et al*., 2009*a*; Geyer *et al*., 2013). Our colleagues have used 3D reaction-diffusion models to simulate these pH_S_ and pH_i_ changes as CO_2_ diffuses into a spherical cell (Somersalo *et al*., 2012; Musa-Aziz *et al*., 2014*a*, 2014*b*; Occhipinti *et al*., 2014; Calvetti *et al*., 2020).

Figure 2*b* shows a representative pH_S_ recording from a H_2_O-injected or “control” oocyte during such a solution change (for solution composition, see Table 1). The vertical gray bands in the figure indicate periods during which we withdraw the tip of the extracellular pH microelectrode from its previous position, in which its tip created a small dimple (∼40 μm) in the oocyte surface, to a distance of ∼300 μm for recalibration in the bECF, which we assume to have a pH of precisely 7.50. We describe the recalibration in the legend of Figure 2.

Figure 2*C* shows a representative western blot of membrane preparations (see Methods) of oocytes injected with H_2_O or cRNA encoding either AQP1-WT or AQP1-C189S. The lower band(s) at ∼25 kDa presumably represents unglycosylated or core-glycosylated AQP1 in the endoplasmic reticulum. The upper bands at ∼37 kDa represent mature-glycosylated AQP1, much of which is presumably at the plasma membrane. Assuming that the AQP1 antibody has similar sensitivities for the WT and C189S proteins, we conclude that *Xenopus laevis* oocytes express similar amounts of each.

Figure 3 shows a series of 12 representative pH_S_ recordings, the first of which, in Figure 3A, is a replicate of Figure 2B. We can see that the ΔpH_S_ induced by the CO_2_ influx in this H_2_O-injected control oocyte is much smaller (upward orange arrow in Figure 3A) than in a day-matched oocyte injected with cRNA encoding hAQP1-WT (upward green arrow in Figure 3B).

We define (ΔpH_S_*)_CO2_ as the difference between the CO_2_-induced ΔpH_S_ value of an oocyte expressing a channel protein (upward black arrow in inset of Figure 3B) and the mean ΔpH_S_ for all day-matched control oocytes (e.g., Figure 3A). This difference is a semiquantitative index of the channel-specific CO_2_ flux (Musa-Aziz *et al*., 2009*a*; Somersalo *et al*., 2012; Calvetti *et al*., 2020). The leftmost bar in Figure 4A represents the mean (ΔpH_S_*)_CO2_ value for 34 oocytes expressing AQP1-WT in the absence of inhibitors.

To assess the importance of the monomeric pore, we examined the effect of pCMBS, which, like HgCl_2_, covalently modifies Cys-189 near the extracellular entrance of the monomeric pore and reduces the component of *P*_f_ due to AQP1 (Preston *et al*., 1993), which we define as *P*_f_*. Preston et al showed that HgCl_2_ has no effect on *P*_f_ if oocytes express the C189S mutant of AQP1 (Preston *et al*., 1993). On the basis of pH_i_ measurements, Cooper and Boron (1998) showed that pCMBS significantly reduces the CO_2_ permeability of oocytes expressing AQP1-WT, but that this effect is absent oocytes expressing AQP1-C189S. In the present study, in which we now monitor pH_S_, experiments on individual oocytes confirm that pretreatment with 1 mM pCMBS (see Methods) reduces the CO_2_-induced ΔpH_S_ in an AQP1-WT oocyte (Figure 3E vs. *B*) but not in an oocyte expressing AQP1-C189S (Figure 3F vs. *C*). pCMBS has no effect on a H_2_O-injected oocyte (Figure 3D vs. *A*), for which ΔpH_S_ is already small. We conclude that pCMBS reduces but does not eliminate the ΔpH_S_ due to AQP1 (Figure 3E vs. *D*).

The leftmost bar in Figure 4B represents the mean (ΔpH_S_*)_CO2_ from 22 oocytes expressing AQP1-C189S in the absence of inhibitors. This (ΔpH_S_*)_CO2_ value is very similar to that for oocytes expressing AQP1-WT (leftmost bar in Figure 4A). A comparison of the first and second bars in Figure 4A and B shows that pCMBS reduces (ΔpH_S_*)_CO2_ by somewhat more than half in oocytes expressing AQP1-WT, but has no significant effect in AQP1-C189S oocytes. These data support the hypothesis that, of the CO_2_ that transits AQP1, a major component moves through the same monomeric pores as H_2_O.

While studying 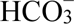 transport, Forster et al unexpectedly found that DIDS blocks a large fraction of CO_2_ permeability of human RBCs (Forster *et al*., 1998). Endeward et al found that, in RBCs genetically deficient in AQP1, DIDS has a reduced effect on CO_2_ permeability (Endeward *et al*., 2006). Here, in the present experiments on single oocytes, we find that pretreatment with 100 μM DIDS reduces the CO_2_-induced ΔpH_S_ in both an AQP1-WT oocyte (Figure 3H vs. *B*) and an AQP1-C189S oocyte (Figure 3I vs. *C*), but not in a H_2_O-injected oocyte (Figure 3G vs. *A*), for which ΔpH_S_ is already small.

The third bars in Figure 4A and B summarize mean (ΔpH_S_*)_CO2_ values from a larger group of oocytes pretreated with DIDS. A comparison of the first and third bars shows that DIDS reduces (ΔpH_S_*)_CO2_ by more than half in oocytes expressing either AQP1-WT (Figure 4A) or AQP1-C189S (Figure 4B). These data are consistent with the hypothesis that a significant fraction of CO_2_ moves through AQP1 by a DIDS-sensitive pathway that is unaffected by the C189S mutation in the extracellular mouth of the monomeric pore.

Returning to experiments on individual oocytes, we see that the sequential exposure of an AQP1-WT oocyte to pCMBS and then DIDS substantially reduces the CO_2_-induced ΔpH_S_ (Figure 3K vs. *B*). Moreover, and the size of the ΔpH_S_ for the pCMBS/DIDS-treated AQP1-WT oocyte (Figure 3K) is about the same as for H_2_O-injected oocytes ±inhibitors (Figure 3A,*D,G,J*). In an AQP1-C189S oocyte, the combination of pretreating with pCMBS then DIDS reduces the CO_2_-induced ΔpH_S_ (Figure 3L vs. *C*), but by no more than DIDS alone (Figure 3I vs. *C*).

The fourth bars in Figure 4A and B summarize mean (ΔpH_S_*)_CO2_ values from a larger group of oocytes, in experiments in which we treated with pCMBS and DIDS in either order. A comparison of the first and fourth bars shows that pCMBS+DIDS reduces (ΔpH_S_*)_CO2_ by slightly more than 100% in AQP1-WT oocytes (Figure 4A). In the AQP1-C189S oocytes, pCMBS+DIDS reduces (ΔpH_S_*)_CO2_ to about the same extent as DIDS alone (Figure 4B), about 40%. Taken together, the data in Figure 3 and Figure 4A and B are consistent with the hypothesis that, in addition to the component of CO_2_ that moves through the four monomeric pores, another component—at least as large—moves through an entirely separate, DIDS-sensitive pathway.

Figure 4*C* summarizes the results of ND96 sham experiments in which—in step F_3_ of Figure 1—we simulated either a 30-min pCMBS exposure (n = 3) or a 60-min DIDS exposure (n = 2). The averages of the two sham groups were nearly identical to that of the control group, showing that the oocytes can tolerate these long protocols.

#### pH_S_ measurements for NH_3_ transport

Figure 5*a* is a schematic representation of the reaction and diffusion events that take place as we exposing an oocyte to a solution containing 0.5 mM NH_3_/NH_4_^+^. Here, the changes in pH_S_ are opposite in direction to those caused by CO_2_/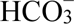exposure. As the weak base NH_3_ enters the cell, it causes a decrease in [NH_3_]_S_, which has two effects. First, it provides a gradient for NH_3_ diffusion from the bECF to the cell surface. By itself, this diffusion, which partially replenishes NH_3_ at the cell surface, has no effect on pH. Second, the decrease in [NH_3_]_S_ also drives the reaction NH_4_^+^ → NH_3_ + H^+^ at the cell surface, which also partially replenishes NH_3_. This reaction produces an acidic pH_S_ transient (Musa-Aziz *et al*., 2009*b*, 2009*a*), so that ΔpH_S_ < 0. Figure 5B shows a representative pH_S_ recording as we introduce NH_3_/NH_4_^+^.

Figure 6 shows a series of 9 representative pH_S_ recordings on individual oocytes, the first of which, in Figure 6A, is a replicate of Figure 5B. We find that the NH_3_-induced ‘−ΔpH_S_’ is much smaller in the H_2_O-injected oocyte (Figure 6A) than in a day-matched oocyte expressing AQP1-WT (Figure 6B). Subtracting the ΔpH_S_ from the H_2_O-injected oocyte from the ΔpH_S_ from the AQP1-WT oocyte yields the channel-dependent, NH_3_-induced ΔpH_S_—(ΔpH_S_*)_NH3_ (Musa-Aziz *et al*., 2009*a*). The leftmost bar in Figure 7A represents the mean (ΔpH_S_*)_NH3_ for 18 oocytes expressing AQP1-WT in the absence of inhibitors.

In experiments on representative oocytes, we find that pretreating an AQP1-WT oocyte with 1 mM pCMBS markedly reduces ‘−ΔpH_S_’ (Figure 6E vs. *B*), but has no effect in an AQP1-C189S oocyte, where ‘–ΔpH_S_’ remains high (Figure 6F vs. *C*); or in a H_2_O oocyte, where ‘–ΔpH_S_’ remains low (Figure 6D vs. *A*). The leftmost bar in Figure 7B represents the mean (ΔpH_S_*)_NH3_ value from 11 oocytes expressing AQP1-C189S in the absence of inhibitors. This (ΔpH_S_*)_NH3_ value is virtually identical to that for oocytes expressing AQP1-WT (leftmost bar in Figure 7A). A comparison of first and second bars shows that pCMBS reduces ‘–(ΔpH_S_*)_NH3_’ by virtually 100% in oocytes expressing AQP1-WT (Figure 7A), but has no effect in AQP1-C189S oocytes (Figure 7B).

Returning to experiments on individual oocytes, we see that DIDS has no effect on ‘−ΔpH_S_’ in a H_2_O-injected control oocyte (Figure 6G vs. *A*), where ‘–ΔpH_S_’ remains low; or in an AQP1-WT oocyte (Figure 6H vs. *B*) or an AQP1-C189S oocyte (Figure 6I vs. *C*), where ‘–ΔpH_S_’ remains high. The third bars in Figure 7A and B summarize mean (ΔpH_S_*)_NH3_ values from larger groups of DIDS-pretreated oocytes. A comparison of the first and third bars in each panel shows that DIDS is without effect on (ΔpH_S_*)_NH3_ in oocytes expressing AQP1-WT (Figure 7A) or AQP1-C189S (Figure 7B). Taken together, the data in Figure 6 and Figure 7 indicate that all of the NH_3_ transiting AQP1 moves via the same pathway as H_2_O and one of the two major components of CO_2_—namely the four pCMBS-sensitive monomeric pores. None of the NH_3_ moves via the alternate, DIDS-sensitive pathway taken by the other major component of CO_2_.

#### Cell-swelling experiments

In parallel with the CO_2_ and NH_3_ assays (Figure 4 and Figure 7), we determined *P*_f_*. In Figure 8A, we summarize our AQP1-WT data, which confirm earlier work that shows that, in oocytes expressing AQP1-WT, mercurials reduce *P*_f_* by about half (Macey, 1984; Preston *et al*., 1993; Zeidel *et al*., 1994; Musa-Aziz *et al*., 2009*a*; Kabutomori *et al*., 2018) but that DIDS is without effect (Macey, 1984; Endeward *et al*., 2006). Figure 8B confirms that the AQP1-C189S mutation renders *P*_f_* insensitive to pCMBS (Preston *et al*., 1993; Cooper & Boron, 1998; Kabutomori *et al*., 2018), and also shows that the mutation does not affect the lack of DIDS sensitivity.

### Modification of hAQP1 by DIDS

Via rapid and reversible electrostatic interactions, the two sulfonate groups of DIDS can reversibly interact with cationic sites on proteins. Via slower covalent reactions with the –NH_2_ group of lysine or –OH/–SH groups of other amino acids, the two isothiocyanate groups of DIDS can act as a homobifunctional crosslinking reagent (Cabantchik & Rothstein, 1972; Lepke *et al*., 1976).

#### pH_S_ experiments on oocytes

To assess whether the DIDS inhibition occurs via an electrostatic or covalent interaction, we followed a DIDS exposure with an albumin wash (see Methods) to scavenge non-covalently-bound DIDS. Figure 9A shows that the albumin wash does not significantly reduce the degree to which the DIDS pretreatment reduces decreases (ΔpH_S_*)_CO_. Because the inhibition is irreversible, the interaction of the DIDS with hAQP1 is most likely covalent. To investigate this covalent modification, we performed the biochemistry experiments (presented in next section) in which we FLAG-tagged hAQP1 at its N-terminus, overexpressed the construct in *Pichia pastoris,* and prepared spheroplasts. Here, in control oocyte experiments, we observe that the FLAG tag has no effect on the (ΔpH_S_*)_CO_ ±DIDS or ±albumin (Figure 9B vs. *A*). We also find that the FLAG tag is without effect in *P*_f_ assays, ±pCMBS or ±DIDS (Figure 9D vs. *C*).

#### Biochemistry experiments

In separate studies, as mentioned in the previous paragraph, we overexpressed FLAG-tagged hAQP1 in *Pichia pastoris*, and prepared spheroplasts. We treated intact spheroplasts with DIDS (or, as shams, without DIDS), solubilized the membranes in detergent, and purified hAQP1 using an anti-FLAG resin. Further purification by size-exclusion chromatography (Figure 10*a*) shows that DIDS treatment promotes the formation of higher– molecular-weight hAQP1 species, as indicated by the increased peak height of the void volume (V_0_). We then pooled and concentrated fractions from each peak. Optical spectroscopy of this material (Figure 10*b*) reveals the expected increase in absorbance centered at ∼340 nm (brown arrow; see Kodippili *et al*., 2009), due to DIDS in the DIDS-treated vs. sham samples. We also separated the pooled/concentrated material by SDS-PAGE, and transferred to membranes for western blotting. In the anti-FLAG blot (Figure 10*C*), the dominant species are hAQP1 monomers (≤50 kDa, arrows a–c) that, as we will see, represent mainly intracellular protein (i.e., not on plasma membrane). Presumed dimers (arrow d) and tetramers (arrow e) are visible only in DIDS-treated samples (lanes 2, 4, 6), and represent proteins accessible to DIDS at the outer cell surface. In the anti-DIDS blot (Figure 10*D*), the monomeric species (dominant in the anti-FLAG blot) are relatively poorly represented, whereas the presumed dimers and tetramers are clearly visible, though only for DIDS-treated samples (lanes 2, 4, 6). These data are consistent with the hypothesis that, once the divalent DIDS molecule reacts with one AQP1 monomer, the odds are high that the DIDS crosslinks to one more to yield dimers and tetramers. Although we attempted to use mass spectrometry to identify AQP1 residue(s) derivatized by DIDS, we were unable to achieve coverage of protein fragments containing predicted DIDS-reactive sites near hydrophobic transmembrane segments.

**Figure 10:**
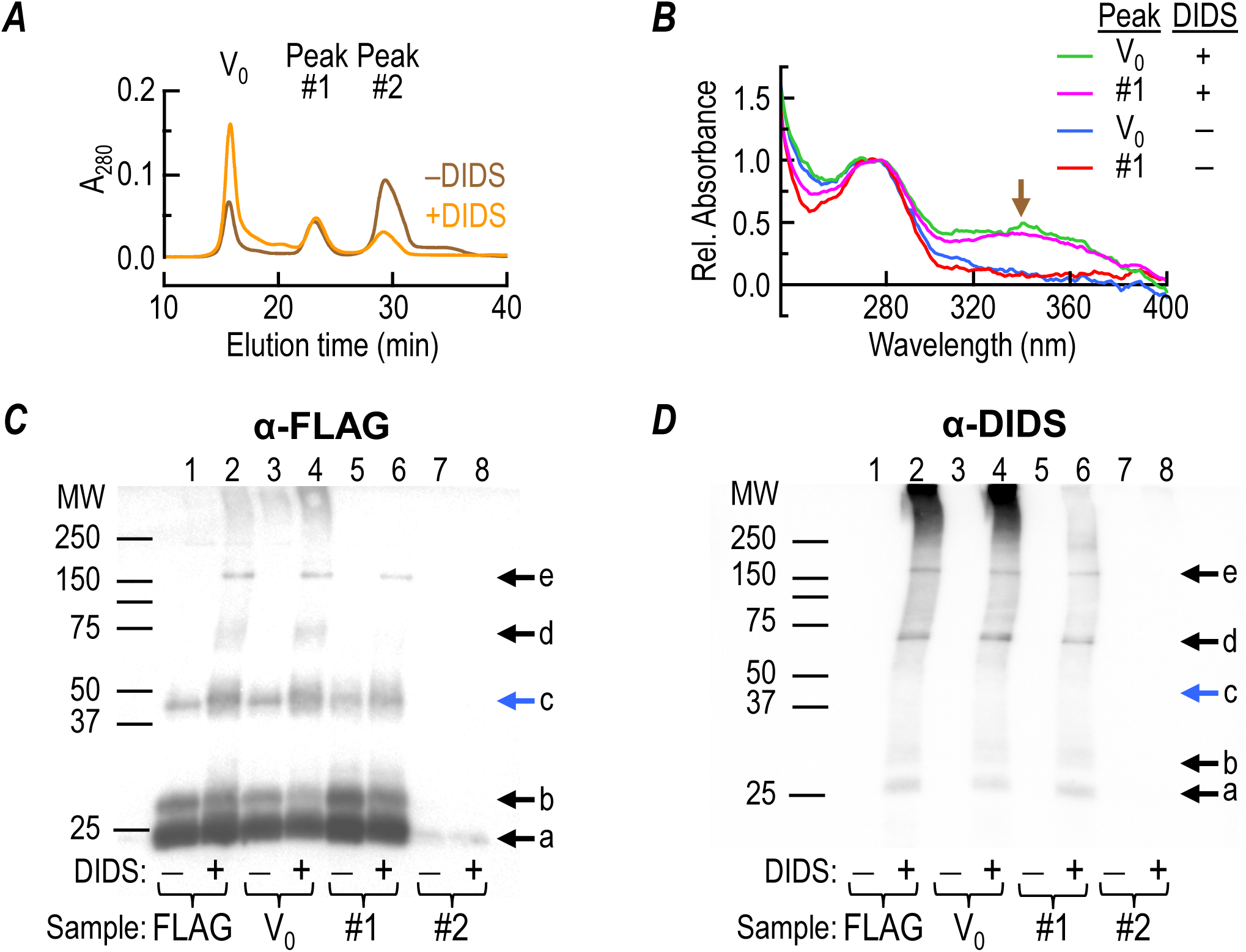
Reaction of DIDS with N-terminally FLAG-tagged hAQP1 overexpressed in *Pichia pastoris*. *A*, Size-exclusion chromatography of solubilized proteins. We treated spheroplasts with DIDS (or no DIDS in parallel, sham experiments), solubilized with DMM, purified N-terminally FLAG-tagged hAQP1-WT using an anti-FLAG column, and then separated by FPLC on a Superdex-200 column, recording A_280_ vs. time. The orange record represents a DIDS-treated sample and the brown record, a control sample (–DIDS) from the same preparation. The first peaks are the void volume (V_0_), which consists of high molecular-weight (MW) proteins. Peak #1 consists of lower-MW proteins, and Peak #2, even smaller ones. We translated both records to so that A_280_ averaged zero between 10 and 13 min, and then we scaled the orange +DIDS record to that it has the same total area under the curve as Peak #1 in the –DIDS record. *B*, Absorbance spectra of material (±DIDS) from peaks V_0_ and #1, from a preparation similar to that shown in panel *A*. We normalized all spectra to unity at 280 nm. The increased absorbance from ∼320 to ∼370 nm in the DIDS-treated samples presumably represents DIDS, which has an absorbance peak at ∼340 nm (Kodippili et al., 2009). C, Western blot of material from the same experiment as in panel *B*, probed with anti-FLAG: Lane 1. Material obtained from spheroplasts not treated with DIDS, and purified on anti-FLAG column, as described for panel *A*, but not subjected to size-exclusion chromatography. ||| Lane 2. Same as Lane 1, but from DIDS-treated spheroplasts ||| Lane 3. Peak V_0_ from spheroplasts – DIDS ||| Lane 4. Same as Lane 3, but from spheroplasts +DIDS ||| Lane 5. Peak #1 from spheroplasts –DIDS ||| Lane 6. Same as Lane 5, but from spheroplasts +DIDS ||| Lane 7. Peak #2 from spheroplasts –DIDS ||| Lane 8. Same as Lane 7, but from spheroplasts +DIDS. The presumed assignments are: “a” band < 25 kDa, unglycosylated AQP1 monomer cleaved in the C-terminus; band “b”, 28 kDa, full-length unglycosylated monomer; band “c”, glycosylated monomer (blue arrow); band “d”, glycosylated dimer; band “e”, glycosylated tetramer. *D*, Western blot of material from the same experiment as in panels *B* and *C*, probed with anti-DIDS. The lane assignments are the same as in panel *C*. The results in panels *A* through *D* are representative of two independent experiments.

## Discussion

### Pathways for NH3 vs. CO2

#### NH_3_ permeation through monomeric pores

Although previous work had shown that AQP1 can serve as a conduit for NH_3_ (Nakhoul *et al*., 2001), the present work is the first direct demonstration that the four monomeric pores of AQP1—which are responsible, of course, for all H_2_O conductance (Preston *et al*., 1993)—are, in fact, also responsible for all NH_3_ conductance. This result is not unexpected, given the hydrophilicity of the monomeric pore (Walz *et al*., 1997; Sui *et al*., 2001; Tajkhorshid *et al*., 2002), and the similarities of the electronic structures of the NH_3_ molecule (i.e., 1 lone pair of electrons + 3 N–H bonds among 4 *sp*^3^ hybrid orbitals) and the H_2_O molecule (i.e., 2 lone pairs + 2 O–H bonds among 4 *sp*^3^ hybrid orbitals). The primary evidence for our conclusion is that pCMBS reduces (ΔpH_S_*)_NH3_ to zero (Figure 7A, center vs. left bars), an effect abrogated by the C189S mutation (Figure 7B, analogous bars). On the other hand, none of the NH_3_ moves via the parallel DIDS-sensitive pathway, inasmuch as DIDS has no effect on (ΔpH_S_*)_NH3_ (Figure 7A and B, right vs. left bars). These results for NH_3_ permeability parallel those for osmotic water permeability in Figure 8.

#### CO_2_ permeation through monomeric pores

The present work confirms earlier observations (Cooper & Boron, 1998) that at least some CO_2_ moves through AQP1 via the monomeric pores and, furthermore, shows that this pathway amounts to about half the total (ΔpH_S_*)_CO2_ signal. This result is not unexpected, given the amphiphilic nature of CO_2_. Moreover, the molecular dynamics simulations of Wang *et al*. (2007) suggest that 40% of the CO_2_ permeability of AQP1 is due to the 4 monomeric pores (i.e., ∼10% per monomeric pore). Supporting evidence for this conclusion in the present study is that pCMBS reduces (ΔpH_S_*)_CO2_ by somewhat more than half (Figure 4A, second bar from left vs. leftmost bar), an effect abrogated by the C189S mutation (Figure 4B, analogous bars).

#### CO_2_ permeation through a parallel pathway

The present work provides the first physiological evidence for a second pathway for CO_2_ through any AQP tetramer. This alternative pathway is apparently parallel to the four monomeric pores. The primary evidence is that (a) DIDS reduces (ΔpH_S_*)_CO2_ by somewhat more than half in hAQP1-WT (Figure 4A, third bar from the left vs. leftmost bar), (b) the C189S mutation does not affect the DIDS blockade (Figure 4B, analogous bars), and (c) the combination of pCMBS and DIDS reduces (ΔpH_S_*)_CO2_ to zero for oocytes expressing AQP1-WT (Figure 4A, rightmost vs. leftmost bars), but still somewhat more than half for oocytes expressing AQP1-C189S (Figure 4B, analogous bars).

### Effect of Inhibitors

#### DIDS

Because the DIDS blockade is irreversible with both hAQP1-WT (Figure 9A) and hAQP1-FLAG (Figure 9B), DIDS probably interacts covalently with nearly all functional AQP1 tetramers on the oocyte surface. Indeed, with DIDS-treated cells, western blots probed with a DIDS antibody (Figure 10*D*) reveal presumed dimers and tetramers, but virtually no glycosylated monomers (compare Figure 10*C* vs. *D,* blue arrows). Thus, we conclude that one or more bivalent DIDS molecules likely crosslinks two or more monomers of almost all AQP1 tetramers on the cell surface. These observations are consistent with the hypothesis that this crosslinking (observed in biochemical experiments) is essential for blockade (observed in physiological experiments). molecular dynamics simulations of Wang *et al*. (2007) suggest that ∼60% of the CO_2_ flux through AQP1 occurs via the hydrophobic central pore that is largely devoid of H_2_O because it is lined by the sidechains of the following residues (from extracellular to intracellular sides): Val-50, Leu-54, and Leu-58 (contributed by TM2), and Leu-174 and Leu-170 (from TM5). The mobility of a gas like CO_2_ through such a near vacuum is ∼10^4^ higher than in liquid water (see (Boron, 2010; Rumble, 2022). Thus, even though central pores may comprise only a small fraction of total membrane surface area, it is possible that such pores—on the background of a membrane with a low intrinsic CO_2_ permeability in the absence of AQP1 (Boron *et al*., 2011)—could make a significant contribution to overall CO_2_ permeability. We propose that the central pore is the anatomic substrate of the DIDS-sensitive component of CO_2_ permeability of hAQP1-WT.

#### Summing the effects of pCMBS and DIDS

The mathematical simulations of Somersalo *et al*. (2012) suggest that—in the region in which (ΔpH_S_*)_CO2_ is most sensitive to the membrane permeability to CO_2_ (*P*_M,CO2_)—(ΔpH_S_*)_CO2_ should rise linearly with log(ΔpH_S_*)_CO2_ (see their figure 7b). If we assume that pCMBS and DIDS each decrease (ΔpH_S_*)_CO2_ by 50%, then each drug must produce a far greater fractional decrease in *P*_M,CO2_. Following the logic of the mathematical simulations, the blockade of the monomeric pore by pCMBS should be accompanied by some degree of blockade of the parallel CO_2_ pathway (e.g., central pore). However, the converse is apparently not true. That is, the blockade of the parallel pathway by DIDS seems not to produce significant effects on the four monomeric pores inasmuch as DIDS has no effect on either (ΔpH_S_*)_NH3_ (Figure 7) or *P*_f_* (Figure 8). Such an overlapping effect of pCMBS on the monomeric and parallel pathways is not unreasonable inasmuch as the target of pCMBS (i.e., C189) is near the outward-facing NPA motif and the selectivity filter of the monomeric pore, both of which are in close proximity to TM2, which lines the extracellular half of the central pore. We propose that (1) pCMBS blocks CO_2_ and NH_3_ traffic through the monomeric pore and part of the CO_2_ traffic through the parallel pathway, and (2) DIDS blocks (most) CO_2_ traffic through the parallel pathway but no traffic through the monomeric pores.

### Molecular Basis of Gas Selectivity

An important unresolved issue is the molecular basis for gas selectivity by AQPs (Musa-Aziz *et al*., 2009*a*; Geyer *et al*., 2013).

#### CO_2_

The present study indicates that the amphiphilic CO_2_ molecule transits AQP1 via 2 pathways, the four hydrophilic monomeric pores and a parallel pathway (e.g., hydrophobic central pore).

#### NH_3_

Here we show that all NH_3_ transits the four hydrophilic monomeric pores of AQP1; that is, none moves via the parallel pathway (e.g., hydrophobic central pore). Considering the broader family of AQPs, we predict that the hydrophobic parallel pathways (e.g., central pores) are never important routes of NH_3_ conductance, but rather that the characteristics of the monomeric pores determine whether a particular AQP has a relatively high permeability to NH_3_ (AQPs 3, 6, 7, 8, 9), or a relatively low NH_3_ permeability (AQPs 2, 4, 5).

We suggest that, for the AQPs known to be permeable to CO_2_ (AQPs 0, 4, 5, 6, 9), CO_2_ transits some combination of monomeric pores and a parallel, more hydrophobic pathway. For the AQPs with limited CO_2_ conductance (AQPs 2, 3, 7, 8), both the monomeric pores and the parallel pathways presumably have less favorable physico-chemical characteristics. Because more-hydrophobic molecules like O_2_ and N_2_ are less likely to transit through the monomeric pores, their permeabilities through AQPs likely depend exclusively on the physico-chemical characteristics of the parallel, hydrophobic pathway. Thus, the gas-selectivity profile of each AQP likely depends on the physico-chemical nature of each gas and the unique set of physico-chemical properties of the (at least 2) potential pathways through the tetramer.

### Concluding Remarks

Our work establishes the proof of principle for blocking two pathways for CO_2_, at least somewhat independently. Elimination of the monomeric pathway, combined with judicious mutations to the hydrophobic alternate pathway, could enable the design of channels with exquisite selectivity among gases. Such designer channels could improve industrial gas handling on a micro-or nanoscale, provide a simple approach for venting metabolically produced CO_2_ in diving or space flight, provide control of CO_2_ and N_2_ conduction in agriculture, and enable precise O₂ control for medical therapies, synthetic biology, and life-support systems in underwater or space environments.

## Additional information

### Authors’ contributions

R.M-A. contributed to the conception and design of the research; performed oocyte experiments; analyzed and interpreted the oocyte data; prepared figures; drafted and edited the manuscript. R.R.G. contributed to the conception and design of the research related to oocyte *P*_f_ experiments as well as *Pichia pastoris* research; performed all the *Pichia pastoris* experiments; analyzed and interpreted all results collected with *Pichia pastoris*; prepared figures; and drafted the manuscript. S-K. L. validated the cDNA clones for hAQP1 used in the experiments and contributed to the interpretation of pH_S_ experiments. F.J.M. contributed to the design of the research; prepared the figures; analyzed the statistics and interpreted results; drafted and edited the manuscript. W.F.B contributed to the conception and design of the research, and also edited the manuscript.

All authors approved the final version of the manuscript and all qualify for authorship, and all those who qualify for authorship are listed.

## Acknowledgements

We thank Duncan Wong for helpful discussions and computer support; Dale Huffman for engineering assistance and helpful discussions. We thank Steven Torontali for designing and fabricating the oocyte chamber; Gerald Babcock for technical assistance and laboratory management; Morley Schwebel; Charleen Bertolini, Rosalyn Forster, and Lesa Goodman for administrative support. W. F. Boron gratefully acknowledges the support of the Myers/Scarpa endowed chair. R. Musa-Aziz gratefully acknowledges Dr. Mark D. Parker (University at Buffalo: State University of New York) for encouragement and helpful discussions, and gives special thanks in memory of Dr. Gerhard Malnic (University of São Paulo, SP, BR) for his unwavering support and advice.

## Supporting information

N/A

## Data Availability Statement

The data that support the findings of this study are available from the corresponding author upon reasonable request.

## Footnotes

1 We code-named the rabbits, and the antisera they produced, “Ren” and “Stimpy.”

2 In some later experiments, we instead put the oocytes and Ca^2+^-free solution into 50-mL Falcon tubes (catalog # 14-959-49A, TFS) and agitated them using a Tube Rotator (Scientific Equipment Products, Baltimore, MD).

3 We apparatus had this dual-solution arrangement to enable the use of out-of-equilibrium CO_2_/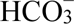 solutions in experiments that are not part of the present study.

4 To obtain *P*_f_ in the proper units (cm s^−1^) in the above equation, the derivation by Zhang *et al*., (1990) requires that the division of “moles of osmotically active particles” (in the Δ*Osm* term) by “moles of water” (in the V_W_ term) yield a numerical result that is unitless, which of course is impossible. This inconsistency arises during the derivation of the water-flux equation when one switches from describing the energy of water (i.e., RT) as “per mole of water” per se to the energy of water “per mole of osmotically active solute” during the introduction of the Van’t Hoff equation. The introduction of a phantom conversion factor with a value of unity—but with the units “(per mole of osmotically active solute)/(per mole of water)”—eliminates the inconsistency.

## Statistics tables

**Statistics Table 4:** Tables of *P*-values for one-way ANOVA with Holm-Bonferroni post-hoc means comparison for comparisons of differences of channel corrected (ΔpH_S_*)_CO_ on exposure of the oocytes to 5% CO_2_/33 mM 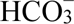. Each table is split in two halves, with FWER α set at 0.05, the upper-right half shows the adjusted α-value for each comparison and the lower-left half the *P*-value. Significant *P*-values are highlighted bold.

| AQP1 |  | Ctrl | +pCMBS | +DIDS | +pCMBS & +DIDS |
| --- | --- | --- | --- | --- | --- |
| | $P \quad \alpha$ | | $\alpha$ | $\alpha$ | $\alpha$ |
| Ctrl |  |  | 0.0100 | 0.0125 | 0.00833 |
| +pCMBS | $P$ | **$7.53 \times 10^{-6}$** | | 0.0500 | 0.0167 |
| +DIDS | $P$ | **$6.46 \times 10^{-5}$** | 0.657 | | 0.0250 |
| +pCMBS & +DIDS | $P$ | **$3.80 \times 10^{-7}$** | 0.0242 | 0.0902 | |

| AQP1-C198S |  | Ctrl | +pCMBS | +DIDS | +pCMBS & +DIDS |
| --- | --- | --- | --- | --- | --- |
| | $P \quad \alpha$ | | $\alpha$ | $\alpha$ | $\alpha$ |
| Ctrl |  |  | 0.0250 | 0.0100 | 0.00833 |
| +pCMBS | $P$ | 0.696 | | 0.0167 | 0.0125 |
| +DIDS | $P$ | **0.00295** | 0.0273 | | 0.0500 |
| +pCMBS & +DIDS | $P$ | **0.00118** | 0.0177 | 0.946 | |

| AQP1 |  | Ctrl | Sham +pCMBS | Sham +DIDS |
| --- | --- | --- | --- | --- |
| | $P \quad \alpha$ | | $\alpha$ | $\alpha$ |
| Ctrl |  |  | 0.0167 | 0.0250 |
| Sham +pCMBS | $P$ | 0.768 | | 0.0500 |
| Sham +DIDS | $P$ | 0.874 | 0.887 | |

**Statistics Table 7.**
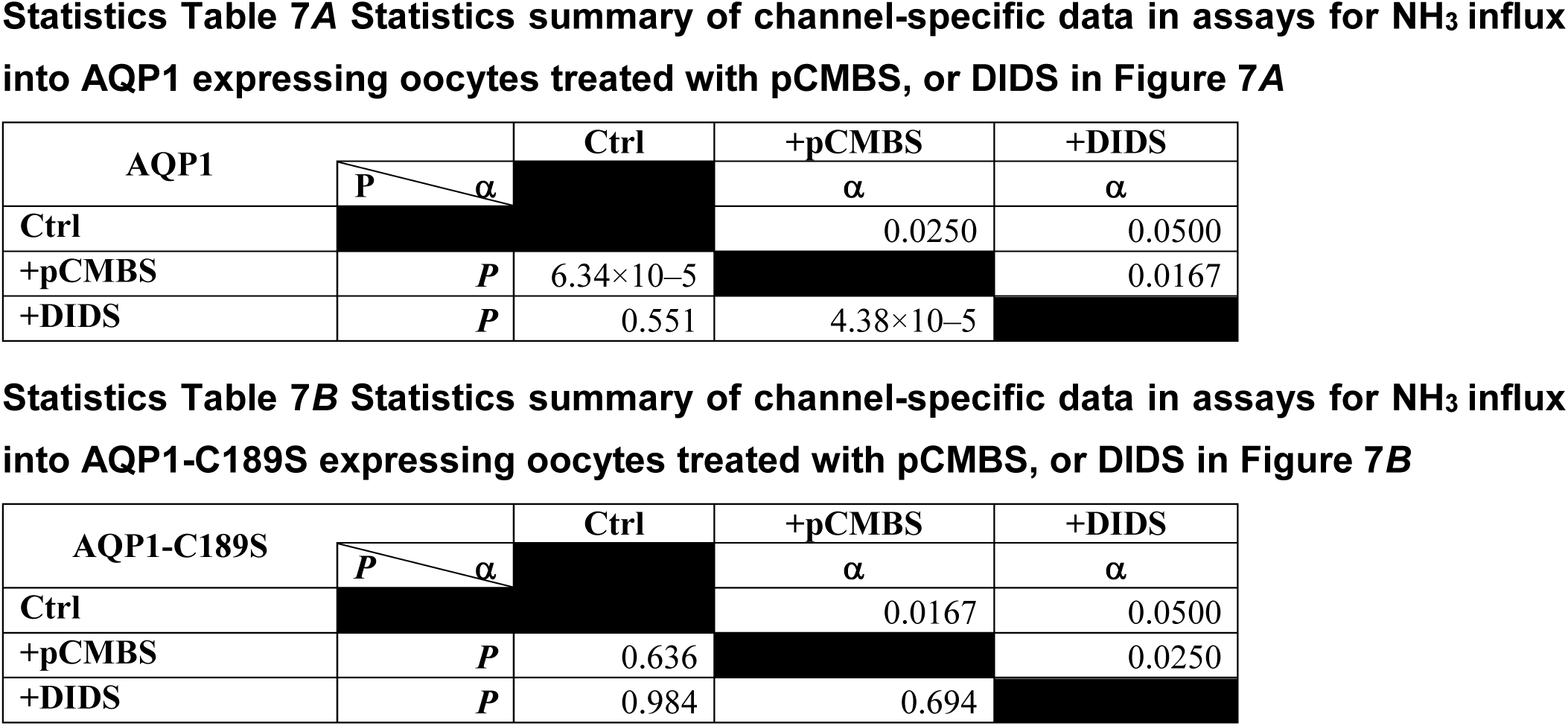
Tables of *P*-values for one-way ANOVA with Holm-Bonferroni post-hoc means comparison for comparisons of differences of channel corrected (ΔpH_S_*)_NH_ on exposure of the oocytes to 0.5 mM NH_4_Cl. Each table is split in two halves by the black-shaded cells, with FWER α set at 0.05, the upper-right half shows the adjusted α-value for each comparison and the lower-left half the *P*-value. Significant *P*-values are highlighted bold.

**Statistics Table 8 for Figure 8.**
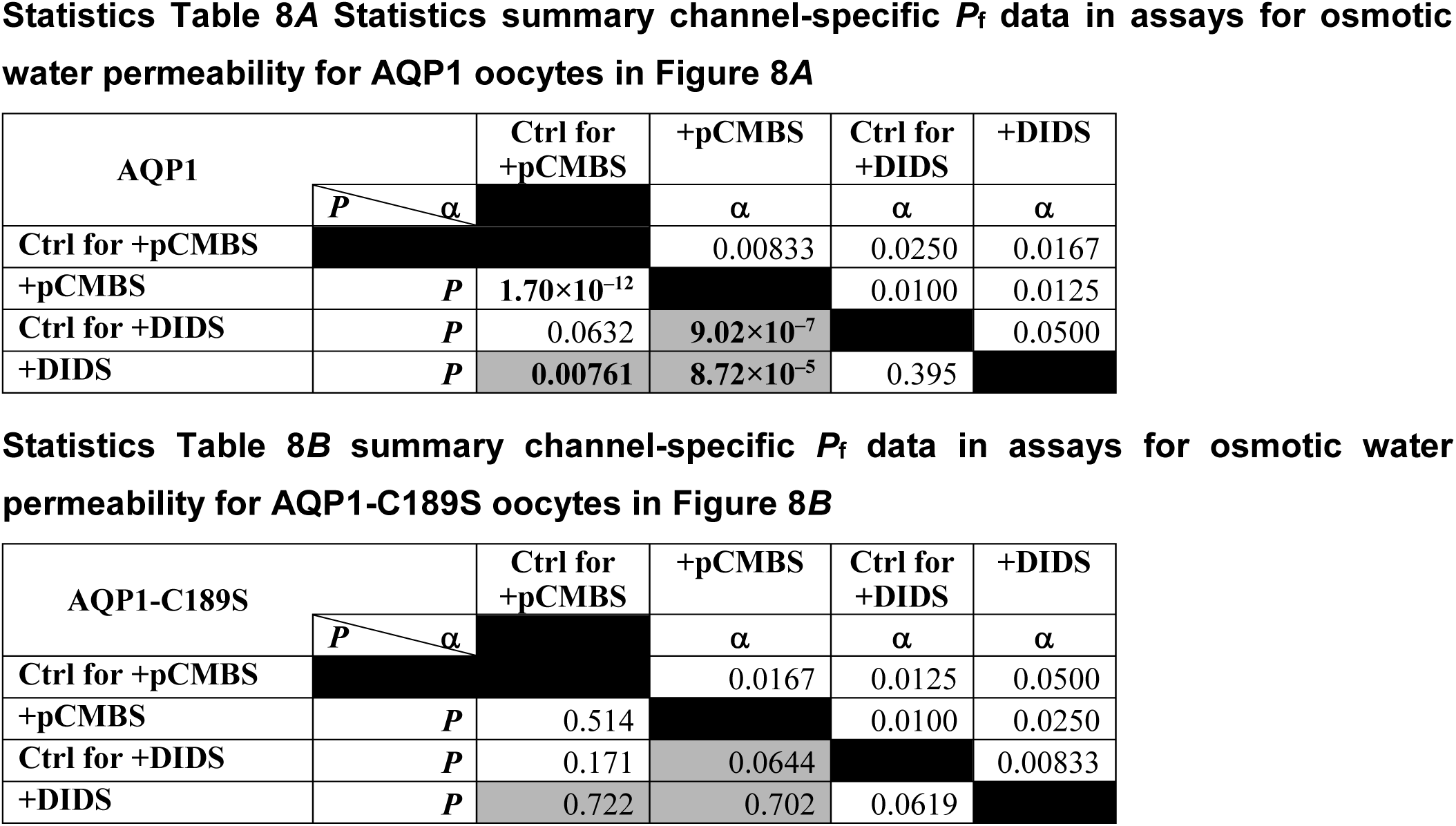
Tables of *P*-values for one-way ANOVA with Holm-Bonferroni post-hoc means comparison for comparisons of differences of channel corrected *P*_f_ * after incubation in no drug control (Ctrl), +pCMBS or +DIDS solutions. Ctrl oocytes are separated into two groups. Those from the same oocyte preparations (frogs) pre-incubated with pCMBS and those from the same oocyte preparations pre-incubated with DIDS. Each table is split in two halves by the black-shaded cells, with FWER α set at 0.05, the upper-right half shows the adjusted α-value for each comparison and the lower-left half the *P*-value. Significant *P*-values are highlighted bold. Cells shaded gray are comparisons between conditions performed on oocytes from different frogs, therefore the comparisons are not pertinent even if the *P*-value is statistically significant.

**Statistics Table 9 for Figure 9.** Tables of *P*-values for one-way ANOVA with Holm-Bonferroni post-hoc means comparison for comparisons of differences in channel corrected *A*, (ΔpH_S_*)_CO_ in WT AQP1 oocytes, *B*, (ΔpH_S_*)_CO_ in AQP1-FLAG tagged oocytes, *C, P* * in WT AQP1 oocytes, and *D, P* * in AQP1-FLAG tagged oocytes. Each table is split in two halves by the black-shaded cells, with FWER α set at 0.05, the upper-right half shows the adjusted α-value for each comparison and the lower-left half the *P*-value. Significant *P*-values are highlighted bold.

**Statistics Table 9A Statistics summary of channel-specific $CO_2$ influx data for into WT AQP1 oocytes**
| AQP1 | $P$ / $\alpha$ | Ctrl | +DIDS | +DIDS +Alb 0.2% |
| --- | --- | --- | --- | --- |
| | | | $\alpha$ | $\alpha$ |
| Ctrl |  |  | 0.0250 | 0.0167 |
| +DIDS | $P$ | **0.00976** | | 0.0500 |
| +DIDS +Alb 0.2% | $P$ | **0.00682** | 0.955 | |

**Statistics Table 9B Statistics summary of channel-specific $CO_2$ influx data for into AQP1-FLAG oocytes**
| AQP1-FLAG | $P$ / $\alpha$ | Ctrl | +DIDS | +DIDS +Alb 0.2% |
| --- | --- | --- | --- | --- |
| | | | $\alpha$ | $\alpha$ |
| Ctrl |  |  | 0.0250 | 0.0167 |
| +DIDS | $P$ | **$3.01 \times 10^{-5}$** | | 0.0500 |
| +DIDS +Alb 0.2% | $P$ | **$4.05 \times 10^{-6}$** | 0.740 | |

**Statistics Table 9C Statistics summary of channel-specific $P_f$ data for into WT oocytes**
| AQP1 | $P$ / $\alpha$ | Ctrl | +pCMBS | +DIDS |
| --- | --- | --- | --- | --- |
| | | | $\alpha$ | $\alpha$ |
| Ctrl |  |  | 0.0167 | 0.0500 |
| +pCMBS | $P$ | **$4.73 \times 10^{-5}$** | | 0.0250 |
| +DIDS | $P$ | 0.361 | **0.00483** | |

**Statistics Table 9D Statistics summary of channel-specific $P_f$ data for into AQP1-FLAG oocytes**
| AQP1-FLAG |  | Ctrl | +pCMBS | +DIDS |
| --- | --- | --- | --- | --- |
| | $P$ $\alpha$ | | $\alpha$ | $\alpha$ |
| Ctrl |  |  | 0.0167 | 0.0500 |
| +pCMBS | $P$ | $7.49 \times 10^{-8}$ | | 0.0250 |
| +DIDS | $P$ | 0.322 | $1.25 \times 10^{-4}$ | |

